# Rapid Validation of AI-designed Drugs via 384-channel Parallel Kinetic Analysis in an Autonomous Laboratory

**DOI:** 10.64898/2026.11.28.769932

**Authors:** Liping Huang, Yuyan Wang, Yuxue Zhang, Yihui Yang, Yuhong Li, Mingqian Chen, Yanan Li, Youqian Chen, Gang Logan Liu

**Affiliations:** School of Food Science and Pharmaceutical Engineering, State Key Laboratory of Microbial Technology, Nanjing Normal University, Nanjing 210023, P. R. China; Zhejiang Collaborative Innovation Center for the Brain Diseases with Integrative Medicine, Zhejiang Key Laboratory of Neuropsychopharmacology, School of Pharmaceutical Sciences, School of Life Sciences, Zhejiang Chinese Medical University, Hangzhou 310053, P. R. China; College of Life Science and Technology, Huazhong University of Science and Technology, 1037 Luoyu Road, Wuhan 430074, P. R. China; The National & Local Joint Engineering Laboratory of Animal Peptide Drug Development, College of Life Sciences, Hunan Normal University, Changsha, Hunan, PR China; Liangzhun Shanghai Industrial Co., Ltd., No. 2467 Tingfeng Highway, Zhujing Town, Jinshan District, Shanghai, China

**Author notes:** These authors contributed equally to this article (Liping Huang, Yuyan Wang, Yuxue Zhang). Corresponding author: (L. Huang), (Y. Chen), (G. L. Liu).

**Keywords:** Nanoplasmonic imaging, AI-driven machine vision, High-throughput kinetics, Unpurified sample analysis, Closed-loop AI drug discovery

## Abstract

While AI-driven protein design rapidly generates vast candidate sequences, experimental affinity screening remains a rate-limiting step in drug discovery. Conventional surface plasmon resonance (SPR) provides real-time kinetics but is limited in channel number, requiring weeks to months for large-scale validation and thereby hindering the design-build-test-learn (DBTL) cycle. Here, we report a fully automated 384-Channel throughput metasurface plasmon resonance (MetaSPR) imaging platform designed to address this throughput challenge. The system integrates three key innovations: a semiconductor-enhanced nanoplasmonic chip providing >3-fold signal amplification, an AI-driven machine vision system enabling truly simultaneous kinetics measurement from all 384 channels at 100 Hz, and a multivalent 3D-hydrogel interface that suppresses non-specific adsorption, enabling direct analysis of unpurified samples. An innovative design combining an automated liquid-handling module with a tubeless parallel detection architecture eliminates sequential scanning, reducing large-scale kinetic screening from weeks to hours. Within a single unattended run, the system completes high-throughput screening of 384 crude samples and a 15 × 15 pairwise epitope binning in 30 minutes, representing a >10-fold efficiency improvement over conventional label-free techniques. To validate its role in the AI loop, we observed conflicting ranking predictions across 141 AI-designed sequences and constructed a 48-candidate adversarial panel to resolve these discrepancies. Screening these binders on our platform (384 kinetic curves in 15 minutes) revealed that physical interface parameters (e.g., ICs) predict experimental K_D_ with far greater accuracy than conventional AI geometric scores (e.g., pLDDT, ipTM). These results indicate that wet-lab kinetic feedback is needed to correct biases in computational scoring. By aligning wet-lab throughput with computational capacity, this platform provides a scalable basis for closed-loop AI-driven biotherapeutic discovery.

## 1 Introduction

Generative protein design has entered an era in which computational output vastly outpaces experimental characterization^1–3^. Fine-tuned diffusion models such as RFdiffusion can now design de novo antibody variable domains and miniprotein binders against user-specified epitopes with atomic-level precision, with cryo-EM validation confirming designed CDR loop conformations^4, 5^. Protein language model–based predictors such as IgFold achieve near-experimental structural accuracy in under 25 seconds, enabling computational interrogation of over one million paired antibody sequences^6^. More recently, AlphaFold 3 extended structure prediction to complexes spanning proteins, nucleic acids, small molecules, and ions, further expanding the scope of computational design^7^. Yet the experimental success of these designs remains limited. De novo designed proteins frequently face intrinsic challenges of aggregation, misfolding, and insolubility that in silico screening does not fully capture; large-scale analyses indicate that only ∼43% of characterized designs meet experimental success criteria and that failure rates vary substantially across topologies^8, 9^. This creates a systemic bottleneck: computational design is now automated, whereas the experimental “test” step has become the sole rate-limiting step of the design-build-test-learn (DBTL) cycle^10^. Without a validation engine that matches the throughput of generation, the cycle does not merely slow down; it stalls.

Existing biosensing technologies cannot fill this gap because each is constrained by an inherent trade-off between kinetic resolution and throughput^11^. Microfluidic SPR instruments, exemplified by the Biacore series, remain the gold standard for real-time kinetic resolution^12–15^. The Biacore 8K operates with eight injection needles and sixteen flow cells, supporting a maximum of eight parallel interactions per run^16, 17^. These systems rely on precisely engineered microfluidic channels that are susceptible to clogging and require highly purified analytes, limiting their compatibility with complex biological matrices^18^. Conversely, ELISA and display-based methods achieve higher throughput but deliver only enrichment scores or semiquantitative endpoint signals. ELISA signals are strongly affected by antibody concentration and cannot resolve affinities in the low-picomolar range^19, 20^. Yeast display enriches binders through wash-dependent sorting; variants with fast off-rates dissociate before detection and are systematically depleted, introducing a kinetic bias before measurement^21^. Array-based SPR imaging platforms such as the Carterra LSA allow 384-ligand printing, but detection is constrained by 48-channel microfluidic flow cells, so the fluidic paths are shared rather than independently addressable across all 384 channels^22–24^. Furthermore, their reliance on microfluidic channels makes them poorly suited to unpurified complex matrices such as crude cell culture supernatants^25^. The field therefore lacks a technology that simultaneously achieves high throughput, high kinetic resolution, and direct compatibility with crude samples.

To bridge the gap between AI-driven molecular design (Fig. 1a) and fully automated wet-lab execution (Fig. 1b), we developed an integrated, automated 384-throughput metasurface plasmon resonance (MetaSPR) imaging platform that rebuilds the sensing architecture along three levels. At the physical level, a CdS-enhanced nanocup array exploits exciton-plasmon coupling to amplify the local electromagnetic field, providing the signal-to-noise ratio required for wide-field imaging across all 384 channels (Fig. 1c)^26^. At the system level, an inverted ultra-large field-of-view transmission imaging system coupled with an encoder-decoder convolutional neural network (CNN) enables automated well recognition and spectral unmixing, extracting real-time binding curves from all 384 channels simultaneously, a parallel imaging modality fundamentally different from the sequential scanning architecture of current array-based instruments^27^. At the chemical level, we engineered a multivalent 3D-hydrogel interface for site-specific capture of His-tagged proteins. Tris-NTA chelation raises the binding affinity by roughly 700-fold relative to mono-NTA (∼10 μM to ∼10 nM) and orients captured ligands more uniformly. Combined with a photo-crosslinked carboxymethyl chitosan hydrogel, this interface suppresses non-specific adsorption even in crude samples and supports repeated chip regeneration. Featuring an integrated liquid-handling module, the platform operates as an automated kinetic screening engine rather than a standalone sensor (Fig. 1b). The AI machine-vision system extracts kinetic curves from all 384 channels and feeds these data back to the design module, establishing a closed-loop feedback mechanism for iterative optimization (Fig. 1d). We propose that this architecture serves as a wet-lab validation layer for AI-driven molecular design, so that experimental throughput need not limit the pace of computational innovation.

**Fig. 1.**
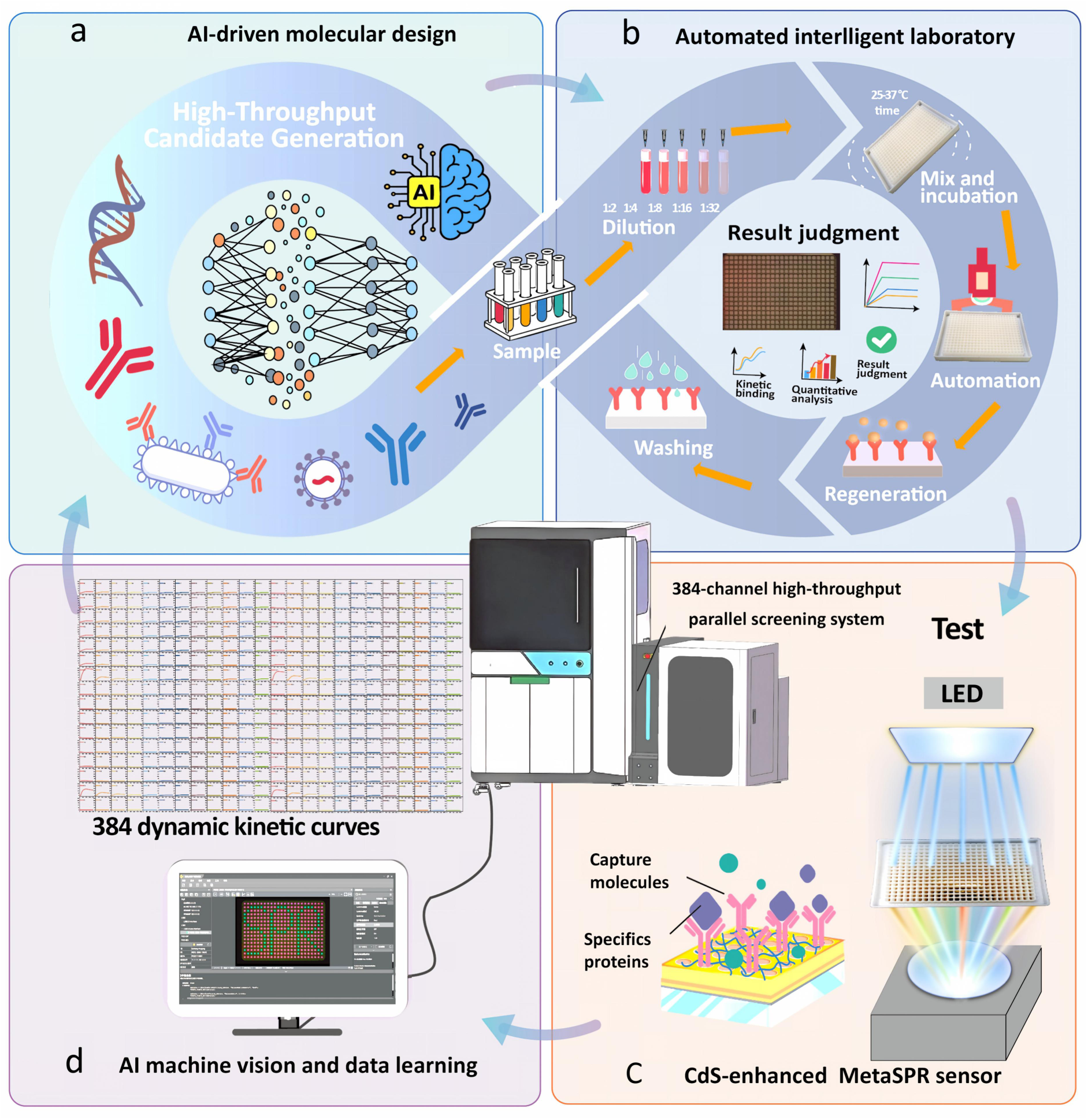
Architecture of the closed-loop MetaSPR high-throughput sensing platform integrating AI-driven molecular design and an automated intelligent laboratory. (a) AI-driven generation of high-throughput candidate biomolecules targeting specific epitopes. (b) Automated intelligent laboratory workflow, encompassing sample dilution, mixing/incubation, real-time sensing, chip regeneration, and automated data interpretation. (c) CdS-enhanced MetaSPR sensor for label-free, high-throughput molecular interaction analysis based on an ultra-large field-of-view imaging system. (d) AI-based machine vision extracting 384-throughput real-time kinetic curves and data learning, establishing a closed-loop feedback mechanism for molecular design.

## 2 Results

### 2.1 Design, Simulation, and Characterization of the CdS-Enhanced Nanocup Array MetaSPR Chip

To achieve 384-throughput high-density wide-field imaging, the sensor chip must maintain a high signal-to-noise ratio (SNR) across the entire imaging area. Conventional Au-only SPR chips provide limited plasmonic field enhancement and are therefore unsuitable for high-throughput readout^28^. To address this, we introduced a semiconductor interlayer between the gold films to amplify the local electromagnetic field. Cadmium sulfide (CdS) was selected for this interlayer because it combines a direct bandgap of approximately 2.42 eV, strong visible-light absorption (∼10^4^ cm^−1^), and pronounced photoconductive properties. This combination allows it to efficiently absorb excitation light and participate in plasmonic field enhancement at the metal-semiconductor interface^29^.

To optimize the MetaSPR chip design, we constructed a dielectric response model (Fig. 2a) and evaluated the electromagnetic field characteristics of two configurations with 20 nm and 40 nm top Au layers using 3D-FDTD simulations. The simulations revealed strong electric field confinement at the nanopillar tips for both structures (Fig. 2b and Supplementary Fig. S1a–b). When the ambient refractive index shifted from 1.3330 to 1.3622, both structures showed a characteristic red shift in their transmission and reflection spectra (Fig. 2c and Supplementary Fig. S1c). The 40 nm Au structure showed the largest peak-to-valley differential responses at approximately 675 and 785 nm (Fig. 2d,e and Supplementary Fig. S1d,e). Its differential signal intensity was markedly higher than that of the 20 nm Au structure, identifying the 40 nm Au top layer as optimal.

**Fig. 2.**
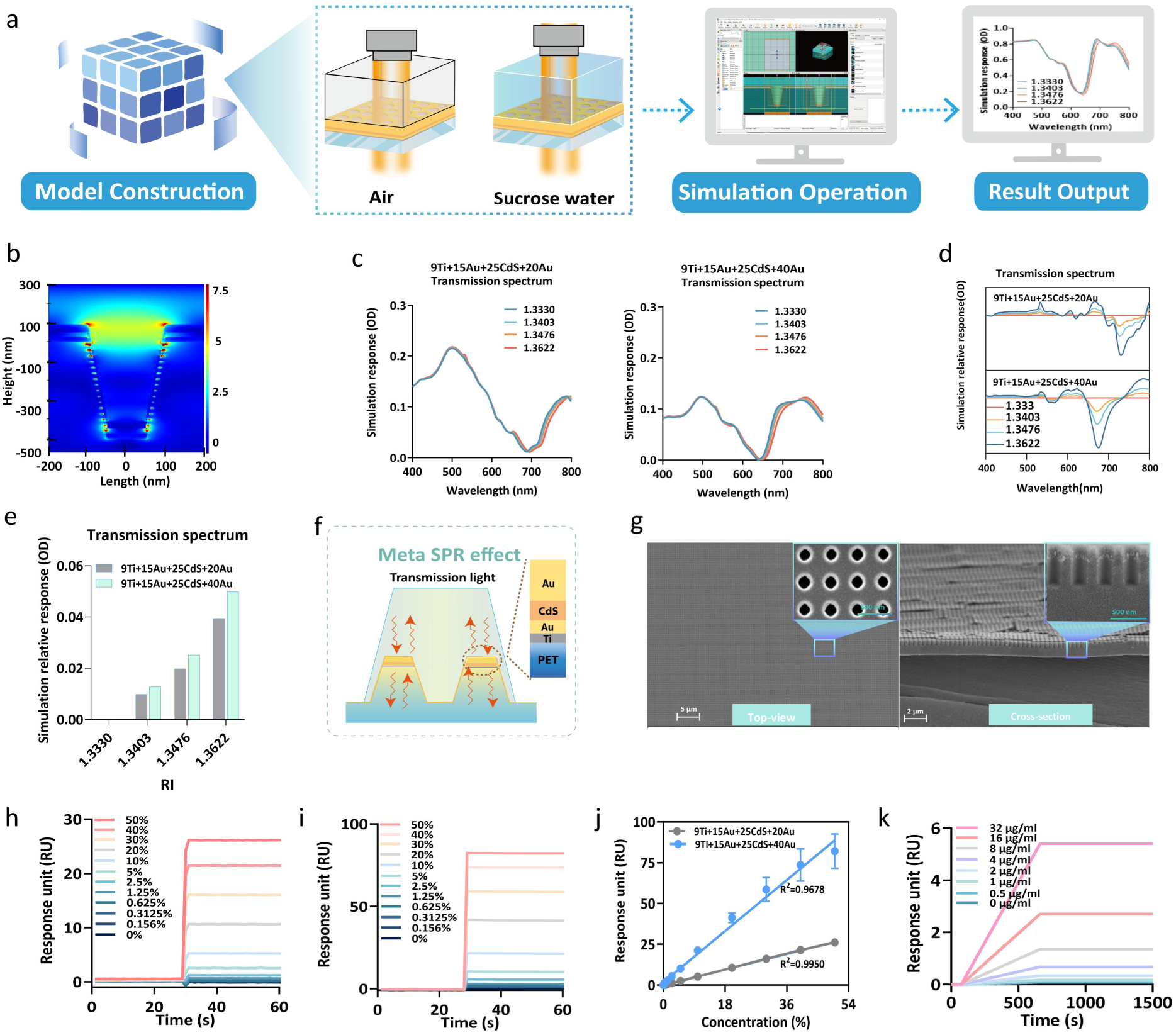
Design, simulation, and characterization of the CdS-enhanced MetaSPR chip. (a) Workflow of the 3D-FDTD simulation. (b) Simulated electric field distribution of the 40 nm Au structure at the optimal resonance wavelength. (c, d) Simulated transmission spectra (c) and normalized spectra (d) for the 20 nm and 40 nm Au structures at varying refractive indices. (e) Comparison of differential SPR signal intensities (peak-to-valley) between the two configurations. (f) Working principle of the CdS-based MetaSPR sensor. (g) SEM image of the fabricated nanocup array. (h, i) Real-time response curves of the 20 nm (h) and 40 nm (i) Au chips toward a sucrose concentration gradient. (j) Linear correlation between the response signals and sucrose concentration. (k) Real-time kinetic sensorgram of Protein G-IgG binding measured on the 40 nm Au chip.

Scanning electron microscopy (SEM) confirmed the high uniformity and morphological consistency of the fabricated CdS nanocup arrays (Fig. 2g). To validate the simulation accuracy, we synchronously recorded the real-time responses to varying sucrose concentrations using the integrated platform (Fig. 2h,i). Across all 96 wells, the 40 nm Au chip produced an endpoint response approximately threefold higher than the 20 nm Au chip (Supplementary Fig. S1f,g).

The limit of detection was 0.156% (w/v), a fourfold improvement over 0.625% for the 20 nm chip, and the response remained linear across the tested concentration range (Fig. 2j).

To further validate the capability of the chip for detecting real biomolecular interactions, we immobilized Protein G on the surface and measured the binding kinetics of IgG at varying concentrations. The 40 nm Au chip clearly resolved concentration-dependent association and dissociation curves (Fig. 2k), whereas the 20 nm Au chip failed to produce detectable signals under identical conditions (Supplementary Fig. S1h). These results show that the CdS-enhanced 40 nm Au structure provides the sensitivity required for 384-throughput kinetic imaging and was therefore adopted as the substrate in all subsequent experiments.

### 2.2 System Architecture and Performance Validation of the Automated 384-throughput High-Throughput Analysis Platform

To achieve high-throughput, fully automated molecular interaction screening, we developed an industrial-grade, fully automated 384-throughput molecular interaction detection system (Supplementary Fig. S2a, b). This instrument overcomes the limitations of standalone instruments and fragmented hardware by integrating, at the architectural level, a CdS-enhanced MetaSPR chip, an inverted wide-field transmission microscope, an AI-driven image analysis unit, and an automated liquid handling module.

For optical detection and data analysis, an LED light source uniformly illuminates the entire 384-well sensor array. Transmitted light signals are collected through a wide-field objective lens and captured at high frequency by an industrial camera (Fig. 3a). To manage the massive volume of image data from 384 channels, we implemented multispectral image analysis driven by an encoder-decoder convolutional neural network (CNN). The network decomposes the raw images into red, green and blue (RGB) spectral channels. By combining adaptive threshold segmentation with morphological filtering, it mitigates pixel crosstalk in high-density arrays and enables simultaneous extraction of 384 independent sensing signals with high fidelity (Fig. 3b). At its hardware core, the system features a three-axis robotic arm that integrates functional modules such as ultra-wide-field optical detection, automatic plate washing, multichannel pipetting, and 4 °C temperature-controlled storage, all operating within a semi-enclosed, contamination-free environment. Through global scheduling by the underlying software, this fully automated system can execute the entire experimental workflow, including gradient dilution, incubation and mixing, sample loading, data analysis, and multi-cycle chip regeneration, with high precision (Fig. 3c). The system thus removes the dependence of conventional SPR on manual intervention and provides an end-to-end, unattended workflow from stock solution handling to kinetic data generation.

**Fig. 3.**
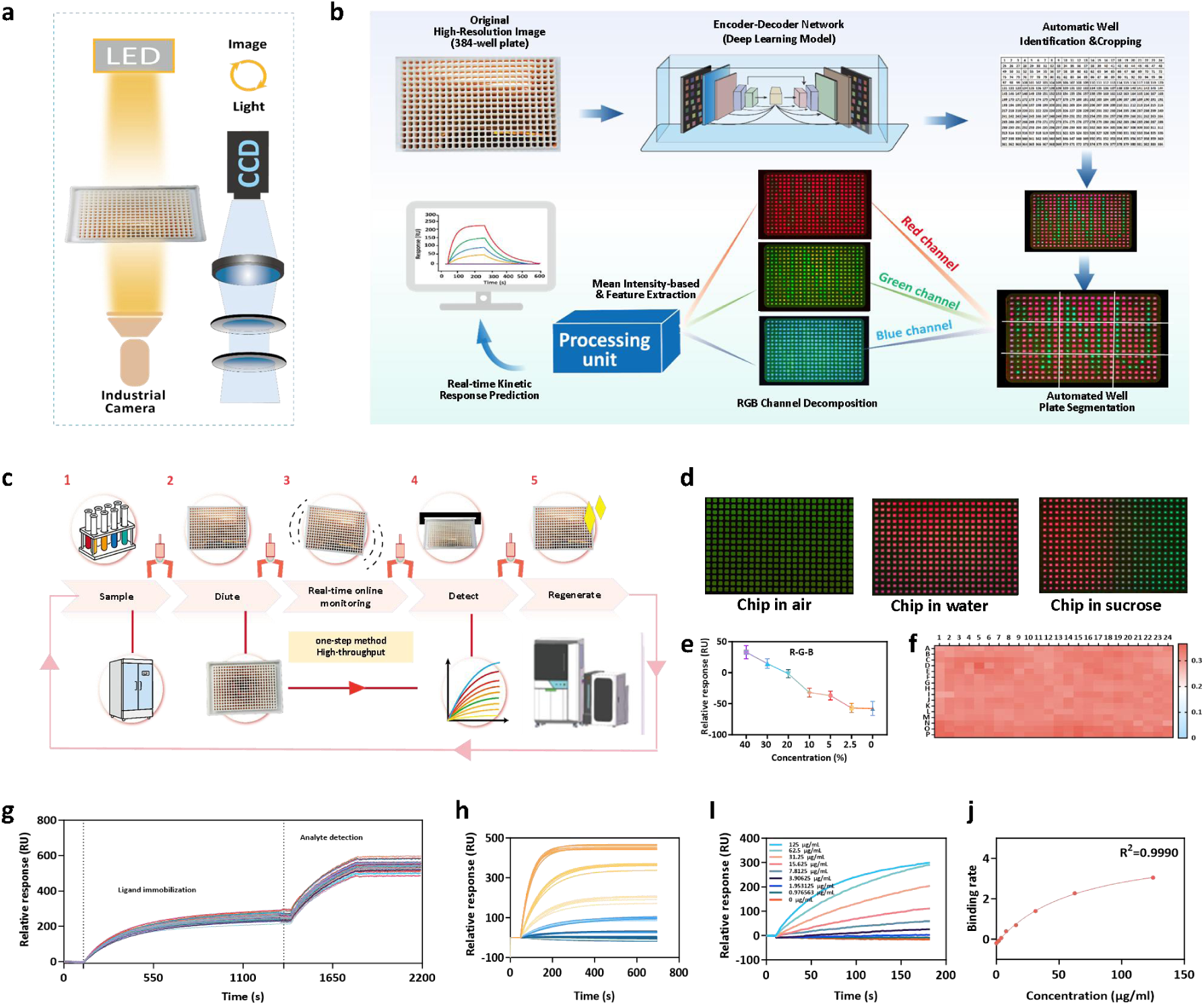
Working principle and basic performance validation of the fully automated 384-throughput high-throughput molecular interaction screening platform. (a) Optical detection: An LED light source uniformly illuminates the entire 384-well sensor array; transmitted light signals are collected by a wide-field objective lens and captured at high frequency by an industrial camera. (b) Schematic diagram of the image acquisition and software data analysis process, including wide-field image capture, multichannel spectral separation, and signal analysis by the processing unit. (c) Complete workflow for fully automated, ultra-high-throughput detection, supporting multiple rounds of continuous cyclic testing. (d) Wide-field imaging results of the 384-well sensor in different media (air, sucrose gradient, water). (e) Consistency validation of parallel detection of sucrose solutions at varying concentrations. (f) Consistency evaluation of 384 independent channels in a single test. (g) Full-process sensing response curve, demonstrating the entire sequence of three consecutive experiments: baseline equilibration, ligand immobilization, and quantitative analyte detection. (h) Sensing curves demonstrating the consistency of IgG antibody binding at different concentrations during high-throughput, multichannel parallel detection. (i) Real-time binding kinetics curves of IgG-protein interactions under different concentration gradients. (j) Results of linear fitting of the initial binding slope signal against analyte concentration, showing excellent linear correlation.

To validate the accuracy of this imaging system, we characterized the light-field response distribution of the chip in air, water and sucrose solutions of varying concentrations (Fig. 3d). Based on the precise decoupling and real-time quantification of multi-channel spectral responses via AI algorithms (Fig. 3e), the system response varied monotonically with the refractive index of the medium. Array-level consistency evaluation showed that the coefficient of variation (CV) across the 384 independent channels was 4.05% in a single test (Fig. 3f), indicating that the AI vision architecture compensates for light-intensity attenuation in wide-field imaging and maintains consistent multichannel readout.

To evaluate the platform’s capture and quantification capabilities in real biological scenarios, we utilized a Protein A (ProA)-modified chip with directionally immobilized antibodies to perform end-to-end real-time kinetic detection of the SARS-CoV-2 RBD protein (Fig. 3g). The complete sensorgram resolved three consecutive stages, baseline equilibration, antibody immobilization and antigen-specific binding, confirming stable anchoring of biomacromolecules at the interface. We then established a rapid quantification module based on the initial binding slope and assessed its quantitative performance in complex matrices. The binding response increased with IgG concentration in a dose-dependent manner, and sensorgrams recorded at the same concentration overlapped closely, confirming channel-to-channel consistency (Fig. 3h). When quantification was based on the binding slope of the analyte during the first 2 min, the system gave a linear response (R^2^ = 0.999) over 0 ∼ 125 μg/mL (Fig. 3i,j). Spike-and-recovery assays showed recoveries between 87.3% and 116.4% across the tested concentration range (Supplementary Table S2). These results support the reliability of the platform for real-time quantification and kinetic analysis and provide a basis for large-scale screening of unpurified crude samples.

### 2.3 Tris-NTA Surface Chemistry Enables Oriented and Reversible Immobilization

To achieve reliable 384-throughput high-throughput screening, the sensing interface must satisfy four criteria: oriented immobilization, high capture capacity, minimal non-specific adsorption, and reversible regeneration. To this end, we integrated Tris-NTA structures into a photo-crosslinked carboxymethyl chitosan matrix to construct a multivalent 3D-hydrogel interface (Fig. 4a), with detailed condition optimization (Supplementary Fig. S3). The trivalent Tris-NTA forms stable hexacoordinate complexes with metal ions (e.g., Ni^2+^, Cu^2+^), enabling site-specific capture of His-tagged proteins while maximally preserving their biological activity^30^. Sucrose gradient tests indicated that this interface maintained high signal intensity while effectively improving spatial uniformity and array consistency (Fig. 4b and Supplementary Fig. S3a). Comparison of the binding responses for three His-tagged model proteins (RBD-V08H, FGFR1, and β2-microglobulin) revealed that the capture signal of the Tris-NTA interface was markedly higher than that of the conventional mono-NTA interface (Fig. 4c).

**Fig. 4.**
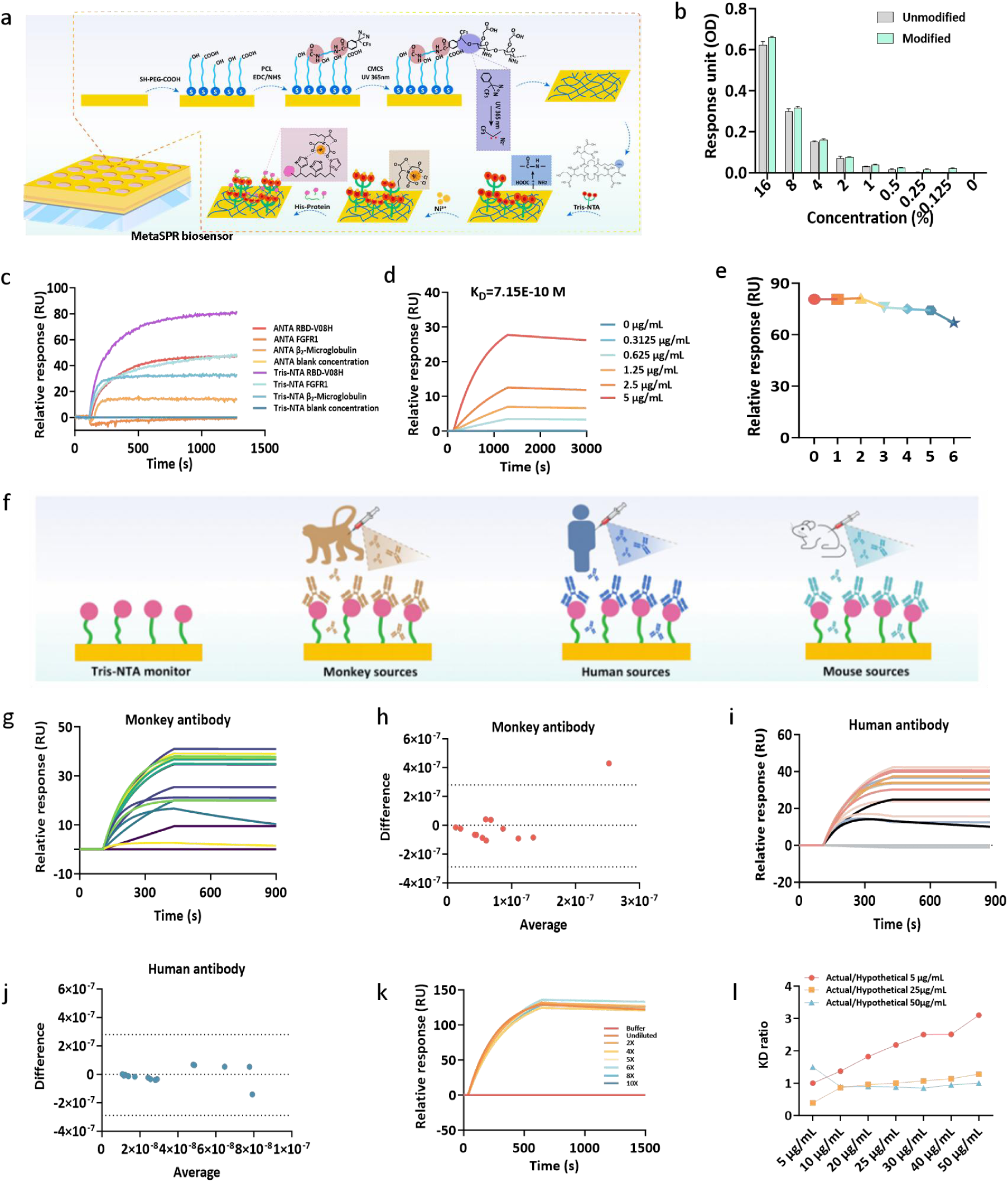
Functionalization, characterization, and optimization of the 3D-hydrogel Tris-NTA-modified chip. (a) Fabrication workflow of the Tris-NTA biosensor: (i) thioglycolic acid self-assembled monolayer (SAM) formation, (ii) EDC/NHS activation, (iii) photo-crosslinker conjugation, (iv) carboxymethyl chitosan hydrogel formation, (v) Tris-NTA coupling, and (vi) Ni^2+^ activation for His-tagged protein capture. (b) Comparison of differential spectra (peak-to-valley) between unmodified and Tris-NTA-modified biosensors at varying sucrose concentrations. (c) Comparison of response values between Tris-NTA and ANTA biosensors toward different His-tagged proteins. (d) Real-time kinetic sensorgram of RBD-V08H binding to its antibody on the Tris-NTA biosensor. (e) Regeneration stability assessment of the Tris-NTA chip. (f) Workflow for detecting antigens from different species using purified antibodies immobilized on the Tris-NTA chip. (g) Real-time kinetic sensorgrams of monkey-derived purified antigens interacting with immobilized antibodies. (h) Consistency comparison of K*_D_* values between the MetaSPR platform and the commercial Biacore 8K system for monkey antigens. (i) Real-time kinetic sensorgrams of human-derived purified antigens interacting with immobilized antibodies. (j) Consistency comparison of K*_D_* values between the MetaSPR platform and the Biacore 8K system for human antigens. (k) Real-time affinity detection of mouse IgG1 samples diluted with PBST to identical final antibody concentrations. (l) Comparison of measured K*_D_* values and the ratios of calculated K*_D_* values based on three different assumed concentrations.

To verify specificity, we monitored the molecular interaction between the SARS-CoV-2 RBD and its specific antibody (S-D001) in real time. The system exhibited a clear dose-dependent kinetic response, yielding an equilibrium dissociation constant (K*_D_*) of 7.15 × 10^-10^ M (Fig. 4d), consistent with previously reported ^31^values; non-specific antibody adsorption was negligible (Supplementary Fig. S3b). After six consecutive immobilization-regeneration cycles, the binding response and kinetic curves for RBD-V08H remained highly consistent (Fig. 4e and Supplementary Fig. S3c), confirming the reversible regeneration capability of the hydrogel interface.

For quantitative accuracy assessment, we performed a parallel comparison using the S-D001/RBD-V08H model against the commercial Sartorius Red384 platform. More than 90% of samples had K*_D_* ratios within a fivefold error window (Supplementary Fig. S4a,b). Subsequently, antibodies from monkey and human sources ( total n = 30) were tested and cross-validated against the Biacore 8K (Fig. 4g-j). Bland-Altman analysis showed that most data points fell within the 95% confidence interval, indicating good agreement between the two platforms.

Conventional SPR has low throughput and requires strict purification, while endpoint methods like ELISA lack kinetic parameters. Cell culture supernatants are compositionally complex and prone to non-specific adsorption. To validate the antifouling capability of this interface, we used Tris-NTA chips to immobilize His-tagged RBD antigens and directly measured the binding kinetics of crude antibody supernatants from different sources (Fig. 4f). To assess matrix effects, crude samples were diluted to a uniform final antibody concentration, which had no significant effect on background responses or K*_D_* values (Fig. 4k and Supplementary Fig. S4c). Across varying dilution factors for samples in the 5 ∼ 50 μg/mL range, the calculated K*_D_* ratios consistently fell within the preset 0.2 ∼ 5.0 window (Fig. 4l). Comparison of sensorgrams before and after regeneration of Tris-NTA chips indicates consistent affinity quantification across regeneration cycles (Supplementary Fig. S4d). These results show that the interface resolves binding kinetics directly in crude supernatants without purification, meeting the requirements of high-throughput automated screening.

### 2.4 384-throughput Parallel High-Throughput Screening of Antibodies in Complex Matrices

Having validated the antifouling and oriented capture capabilities of the 3D-hydrogel Tris-NTA interface, we further evaluated the platform for direct screening of unpurified biological samples. We utilized the Tris-NTA chip to specifically capture His-tagged antigens (SARS-CoV-2 RBD protein), followed by direct injection of diluted crude cell culture supernatants containing target antibodies for real-time kinetic screening (Fig. 5a). Leveraging the simultaneous acquisition capability of the platform, real-time binding signals were automatically extracted from all 384 channels (Fig. 5b). This design prevents weak specific signals from being masked by background interference in complex matrices.

**Fig. 5.**
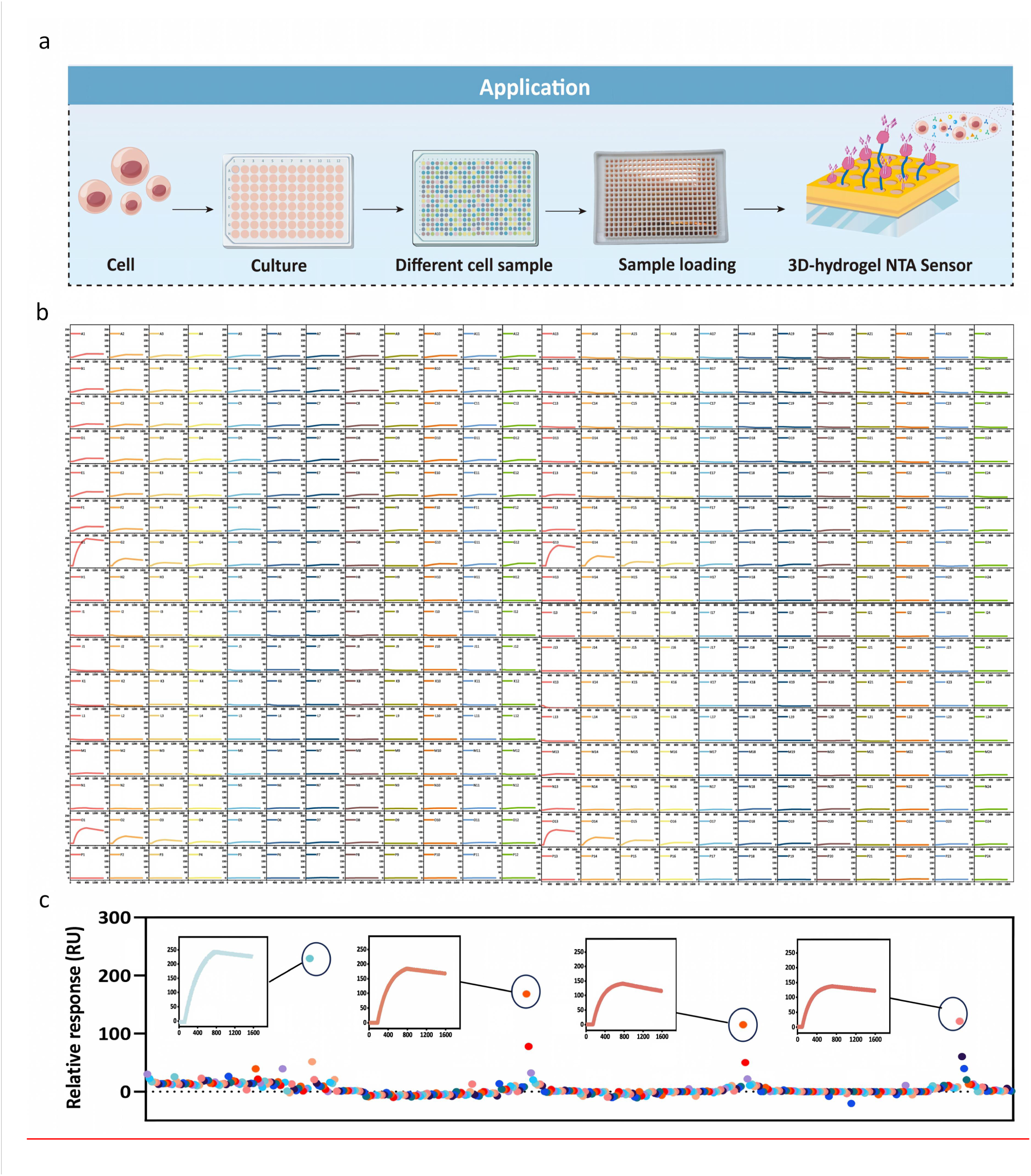
Real-time high-throughput detection of crude cell culture supernatants using the 384-throughput MetaSPR platform. (a) High-throughput screening workflow for crude supernatants based on the 3D-hydrogel Tris-NTA sensor: after oriented capture of His-tagged antigens on the chip surface, diluted crude supernatant samples (containing target antibodies) are directly injected for real-time monitoring without prior purification. (b) Real-time kinetic sensorgrams of crude supernatant samples synchronously acquired from all 384 sensing channels in a single assay. (c) Distribution of endpoint response signals across the 384 channels, along with representative binding kinetics curves corresponding to high-response signals.

To assess intra-assay reproducibility, 96 independent crude supernatant samples were distributed across the 384 channels (four replicates per sample, n = 4). The 15 high-binding candidates identified in the primary screen gave consistent endpoint responses across the four replicates (Fig. 5c and Supplementary Fig. S5a,b), confirming the reproducibility of the platform. In contrast to the stringent purity requirements of conventional SPR and microfluidic systems, the antifouling properties of the Tris-NTA interface enabled direct kinetic analysis of antibody–antigen interactions from crude cell supernatants. This provides a reliable approach for high-throughput antibody screening in complex biological environments and identifies target candidates for subsequent 15×15 epitope binning analysis.

### 2.5 Epitope Binning of Crude Supernatants on the Fully Automated 384-throughput Platform

Epitope-specific antibody identification typically uses animal immunization or antibody library screening followed by epitope mapping, which is laborious, slow and may fail to capture antibodies targeting therapeutically relevant epitopes^32, 33^. Using the continuous unattended operation and antifouling interface of the fully automated 384-throughput system (Supplementary Fig. S6a), we performed high-throughput epitope binning on the 15 high-affinity crude supernatant samples obtained from the primary screen. Epitope binning identifies non-competing antibody pairs that can be combined into therapeutic cocktails or bispecific antibodies. Driven by the automated liquid-handling module, the system completed cycles of gradient dilution, sample loading, detection and chip regeneration without sample purification (Fig. 6a). To minimize background interference from the complex matrix, the optimal dilution of the crude supernatants was determined to be fourfold in sandwich-assay pre-tests (Supplementary Fig. S6b-d).

**Fig. 6.**
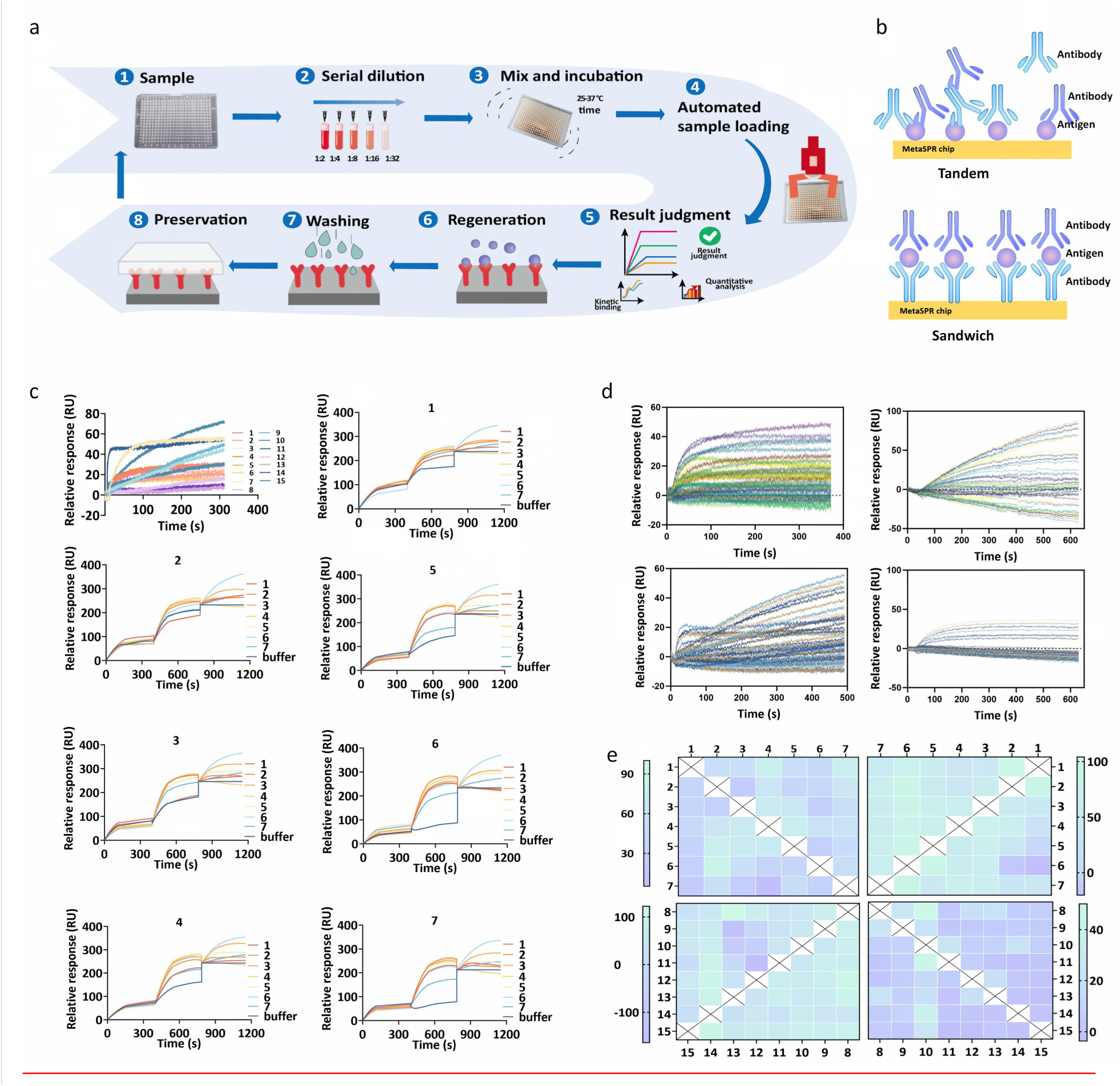
Fully automated 384-throughput platform for epitope binning of crude cell culture supernatants. (a) Fully automated workflow for detection and epitope binning, supporting multiple consecutive cycles. (b) Schematic illustration of the tandem and sandwich binding modes for epitope analysis. (c) Real-time kinetic sensorgrams for the antigen-binding activity of the 15 selected samples, and representative three-stage sensorgrams obtained via the sandwich strategy (sequentially corresponding to primary antibody immobilization, antigen binding, and secondary antibody binding). (d) Real-time kinetic detection results of epitope interactions: tandem assay (left) and sandwich assay (right).(e) Heatmaps of epitope binning results: tandem assay (left) and sandwich assay (right).Samples 1-7 are grouped for epitope pairing, and samples 8-15 are grouped for epitope pairing.

All 15 samples showed clear antigen-binding activity in real time, satisfying the prerequisite for subsequent pairing analysis.We then evaluated these 15 samples using two complementary assay formats, tandem and sandwich assays, and generated a 15×15 matrix combining data from both approaches.(Fig. 6b,c). Owing to the 384-throughput parallel acquisition architecture and tubeless sample introduction design, the full cross-pairing experiment comprising 225 interactions was completed in approximately 30 min of automated operation, with the antigen-modified chips prepared in advance. In contrast, conventional high-throughput bio-layer interferometry (BLI) typically requires >8 h for pairings on the same scale, an order-of-magnitude reduction in assay time^34^.

The epitope-binning heatmaps obtained with the two detection modes revealed epitope differences among the samples (Fig. 6d,e). Several antibody pairs (e.g., Ab2-Ab4, Ab3-Ab4, Ab8-Ab9 and Ab10-Ab14) exhibited clear non-competing signatures, identifying them as candidates for orthogonal pairing. To assess data reliability, Bland-Altman analysis was used to evaluate agreement between the tandem and sandwich results. Most data points fell within the 95% confidence interval, indicating good agreement between the two formats (Supplementary Fig. S6e, Fig. S7 and Fig. S8). Furthermore, the three-stage kinetic sensorgrams obtained via the sandwich strategy clearly resolved the sequential steps of primary antibody immobilization, antigen binding and secondary antibody binding. These results show that the platform performs multi-step epitope binning directly in unpurified matrices, providing an automated workflow for screening and characterization of antibody candidates.

### 2.6 Closing the Loop: High-Throughput Kinetic Validation of AI-Designed PD-L1 Binders

Having completed high-throughput primary screening and epitope binning of crude samples, we extended the platform’s scope to address a central bottleneck in AI-driven drug discovery: bridging the gap between computational design and experimental validation. As generative AI can now efficiently produce massive candidate sequences, accurately identifying those with genuine binding activity has become a rate-limiting step. We first analyzed 141 de novo designed PD-L1 sequences generated by various AI models (Fig. 7a-c and Supplementary Fig. S9). Correlation analysis of computational metrics revealed strong intra-category correlation, for example between the AI confidence metrics pLDDT and ipTM (ρ=0.860), as well as among the physical interface metrics ICs, PD-L1 Contacts, and PRODIGY ΔG (ρ = 0.596 - 0.763). However, the two categories were largely decoupled, with correlations between AI confidence and physical parameters of only ρ=0.169-0.413 (Fig. 7d and Supplementary Fig. S10). This indicates that high confidence in AI structural prediction does not guarantee favorable physical contacts or thermodynamic stability. Furthermore, benchmarking against 15 literature-validated binders (Supplementary Fig. S11) showed that the interface metrics correlated most strongly with experimental affinity (ICs: *ρ*=0.667, contacts: *ρ*=0.609), whereas the predicted binding-energy metrics showed only weak, non-significant trends (PRODIGY ΔG: *ρ*=0.468; PRODIGY KD: *ρ*=0.488 and the AI geometric metrics were not significant (ipTM *ρ*=0.185 and pLDDT *ρ*=0.250). For this benchmark set, interface-derived metrics were thus more informative than either confidence scores or predicted thermodynamic terms. However, because the 141 candidates were ranked very differently by the five core metrics, and single-metric screening is prone to false negatives (Fig. 7e-f), filtering on computational metrics alone carries a high risk of misjudgment.

**Fig. 7.**
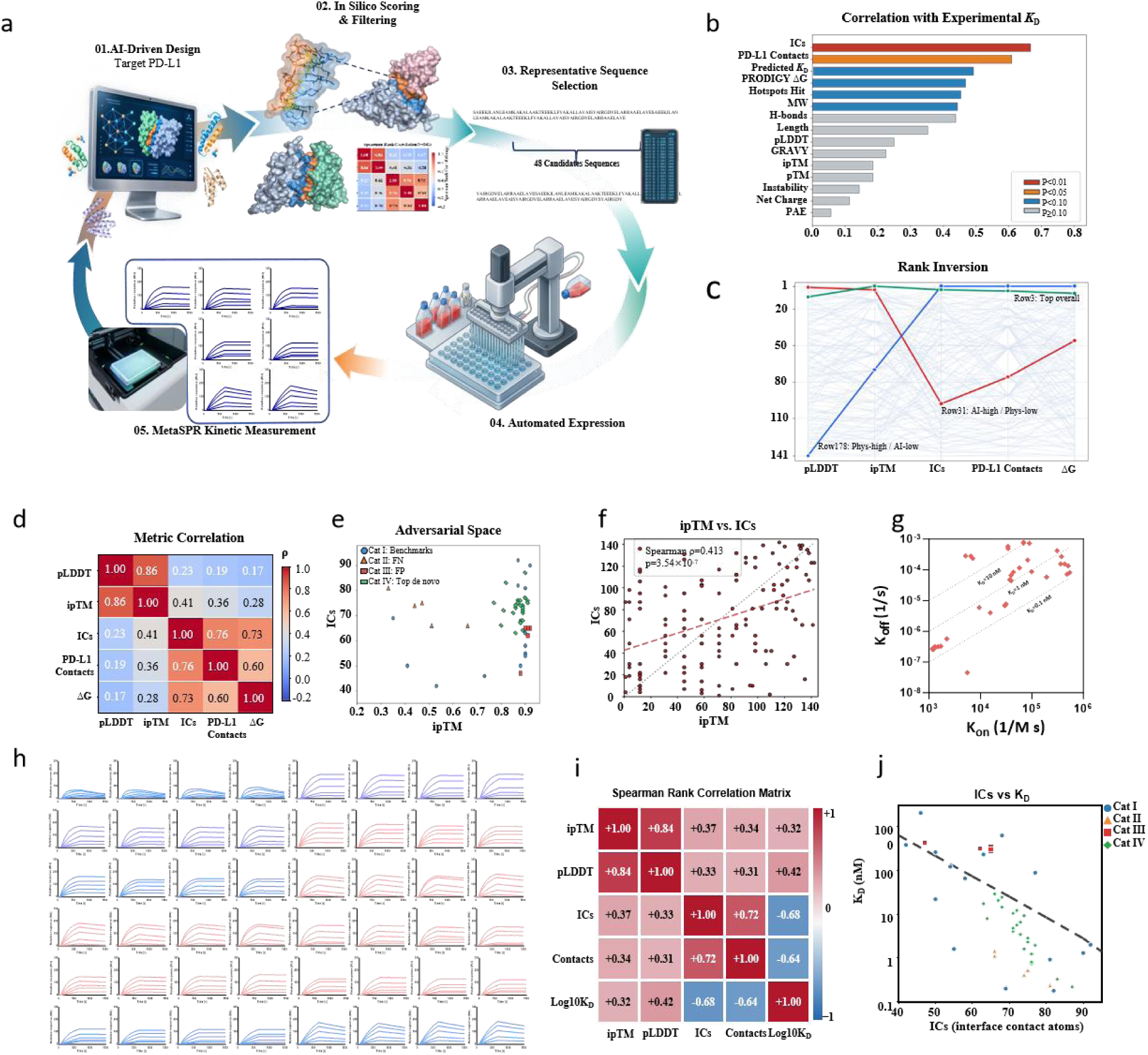
Closing the loop: High-throughput kinetic validation of AI-designed PD-L1 binders. (a) Workflow diagram of the AI-driven wet-dry closed loop. (b) Predictive performance of 15 computational metrics: Spearman correlation coefficients against experimental K*_D_* (n=15). (c) Parallel coordinates plot of 141 candidate molecules across five core metrics (1 = best, 141 = worst). (d) Correlation heatmap of five core metrics (N=141). AI geometric dimension: pLDDT, ipTM; Physical interface dimension: ICs, PD-L1 Contacts, PRODIGY ΔG. (e) Grouping of the adversarial testing panel comprising 48 candidates. Cat I (blue circles): Literature-validated binders (n=15); Cat II (orange triangles): High physical/Low AI (n=5); Cat III (red squares): High AI/Low physical (n=5); Cat IV (green diamonds): Multi-gate de novo designs (n=23). (f) Correlation analysis between ipTM and ICs (Spearman ρ=0.413, p=3.54×10^−7^, slope = 0.39). (g) Scatter distribution of association (k*_on_*) and dissociation (k*_off_*) rate constants derived from 384-throughput kinetic fitting. (h) Real-time kinetic curves of eight concentrations of analyte acquired in parallel across 384 sensing channels. (i) Correlation analysis between the experimental K*_D_* of 48 AI-designed molecules and four core metrics (pLDDT, ipTM, ICs, PD-L1 Contacts). (j) Correlation analysis between experimental K*_D_* and the physical interface metric (ICs).

To quantify the bias of purely computational filtering, we assembled an adversarial test panel of 48 candidates (Supplementary Table S2). The panel contained high-quality de novo designs obtained through multi-gate filtering (Cat IV) together with two types of challenging samples: computational false negatives (binders rejected by the AI filters but validated in the literature, Cat I/II) and AI false positives (designs with high AI geometric scores but insufficient physical contacts, Cat III). Using the 384-throughput MetaSPR platform, we simultaneously evaluated these 48 samples across eight concentrations. A single 15-min run yielded 384 real-time kinetic curves. The distribution of association (k*_on_*) and dissociation (k*_off_*) rate constants obtained by fitting all channels shows that large-scale kinetic parameters can be extracted in parallel from a single experiment (Fig. 7g,h).

The experimental kinetic data revealed a substantial divergence between the experimentally measured affinity ranking and predictions based on AI geometric scores. AI false positives (Cat III), which the AI models ranked highly, bound more weakly than expected. Conversely, computational false negatives (Cat I/II) removed by the computational filters because of low AI scores showed high binding affinities. Correlation analysis showed that measured K*_D_* values correlated strongly with physical interface parameters but only weakly with AI geometric metrics (ipTM, pLDDT) (Fig. 7i,j). This result is consistent with the benchmarking of the 15 literature-validated binders and confirms that static geometric scores poorly reflect actual binding potential. Feeding the kinetic data acquired by the MetaSPR platform back into the design model as a reward function establishes a closed-loop pipeline linking AI design, computational filtering, wet-lab arbitration and model optimization.

## 3. Discussion

The MetaSPR platform described in this work mitigates the inherent trade-off between high kinetic resolution and high throughput that has long constrained molecular interaction analysis. By integrating CdS-enhanced nanoplasmonic sensing, a multivalent 3D-hydrogel interface, AI-driven machine vision, and automated liquid handling, we have established a 384-throughput detection platform capable of real-time, label-free kinetic profiling of both purified and unpurified biological samples. Because AI-driven molecular design continuously generates large candidate sets while high-throughput experimental validation remains a bottleneck, this platform offers a practical means of resolving this conflict.

A key physical mechanism of this platform is the CdS semiconductor interlayer within the metal-dielectric-metal stack. The approximately 3-fold signal enhancement observed experimentally is consistent with the FDTD simulations and arises from two complementary mechanisms: enhanced light absorption and charge separation at the Au-CdS interface, which improves local refractive index sensitivity, and exciton–plasmon coupling, which amplifies the localized electromagnetic field at the nanopillar ^35–39^. This enhancement strategy is not limited to CdS. Other transition metal dichalcogenides (e.g., MoS_2_, WS_2_) and II-VI semiconductors with suitable bandgaps are theoretically predicted to exhibit similar plasmonic enhancement effects, providing a reference for future optimization of sensing materials^40, 41^.

The combination of CdS-enhanced signaling and large-field-of-view imaging is essential for synchronous 384-throughput detection. In conventional SPR imaging, expanding the imaging field inevitably distributes illumination power and reduces the signal-to-noise ratio (SNR) at individual sensing spots^42, 43^. CdS enhancement increases the optical signal per unit area, so that the SNR remains sufficient to extract high-fidelity kinetic curves at 384 densely packed sensing spots. This physical mechanism underpins the high multiplexing capacity of the MetaSPR platform without compromising temporal resolution.

Currently, no existing commercial system simultaneously achieves 384-throughput synchronous imaging, complex matrix compatibility, and fully automated operation. The Carterra LSA-XT offers 384-spot printing but relies on 48-channel microfluidic flow cells and sequential reading, lacking all-channel parallel imaging, which limits its compatibility with unpurified matrices^44, 45^. The Biacore 8K delivers gold-standard kinetic precision but is constrained by an 8-channel parallel fluidic system and stringent sample purity requirements^46^. The Octet BLI platform offers the convenience of a 384-well plate format but lacks the temporal resolution of imaging-based SPR^47, 48^. In this context, the MetaSPR platform occupies a distinct technological niche by combining the throughput of array-based SPR, the kinetic resolution of conventional SPR, and the sample compatibility required for automated workflows.

An important application of this platform is bridging the gap between computational design and experimental characterization. As the PD-L1 case shows, AI models can generate large candidate libraries, but the design-build-test-learn (DBTL) cycle stalls when experimental validation cannot keep pace. By measuring the kinetics of 48 AI-designed binders at eight concentrations directly from crude supernatants (384 data points in total), the MetaSPR platform restores this feedback loop. Moreover, the platform provides kinetic rate constants (k*_on_* and k*_off_*) rather than static affinity values, which is important for training next-generation generative AI models. Coupling this platform directly to AI optimization pipelines could ultimately yield an autonomous closed-loop system that iteratively refines design algorithms using wet-lab kinetic data.

The platform still has several limitations. First, the CdS chip currently relies on electron-beam evaporation, a batch process with inherent wafer-to-wafer variability; to achieve scalable production, a transition to scalable deposition methods such as magnetron sputtering or atomic layer deposition will be necessary^49^. Second, the current refractive index resolution of 2×10^−5^ RIU is sufficient for protein-protein interactions but would need improvement for very weak interactions or small molecules. Third, although 15×15 epitope binning has proven feasible, expanding the matrix to 96×96 or larger will require further optimization of the surface chemistry to maintain uniform ligand density over long experiments.

Beyond antibody discovery, the MetaSPR platform can be broadly applied to various molecular interaction scenarios, including small-molecule candidate profiling, aptamer screening, protein interaction network mapping, and clinical biomarker validation. By combining CdS-enhanced sensitivity, an antifouling interface, 384-throughput real-time imaging and full automation, this platform provides a general platform for data-intensive molecular interaction analysis.

## 4 Conclusions

The MetaSPR platform described here converts kinetic binding measurement from a low-throughput, purification-dependent procedure into a fully automated 384-throughput operation that accepts crude samples directly. Three components make this possible: a CdS-enhanced nanocup chip that supplies the signal-to-noise ratio needed for wide-field imaging, an AI machine-vision pipeline that reads all 384 channels simultaneously, and a multivalent Tris-NTA hydrogel interface that provides oriented capture, antifouling and regeneration. On this basis, primary screening of crude supernatants, 15×15 epitope binning and kinetic profiling of AI-designed binders were each completed within a single unattended run, and the resulting kinetic data exposed systematic errors in purely computational scoring of designed binders. The platform therefore supplies the experimental layer that the design–build–test–learn cycle has lacked: it measures what computational models cannot yet predict, returns these measurements as quantitative constraints for model refinement, and does so at a throughput that matches generative design. Extending chip fabrication to wafer-scale deposition and the interface to larger matrices should make this closed-loop strategy applicable to broader classes of molecular interactions.

## 5. Materials and Methods

### 5.1 Materials and Chemicals

1-Ethyl-3-(3-dimethylaminopropyl)carbodiimide hydrochloride (EDC), N-hydroxysuccinimide (NHS), thioglycolic acid, photo-crosslinker 2, carboxymethyl chitosan, 2-(N-morpholino)ethanesulfonic acid (MES), Na,Na-bis(carboxymethyl)-L-lysine, fish gelatin, sucrose, Tween 20, HEPES, EDTA, imidazole, anhydrous copper chloride (CuCl_2_), nickel chloride (NiCl_2_), cobalt chloride (CoCl_2_), and zinc sulfate (ZnSO_4_) were purchased from commercial suppliers. Sodium hydroxide, hydrochloric acid, sodium chloride (NaCl), potassium chloride (KCl), disodium hydrogen phosphate (Na_2_HPO_4_), and potassium dihydrogen phosphate (KH_2_PO_4_) were of analytical grade. Tris-NTA was synthesized in-house. The model proteins, including β2-microglobulin, FGFR1, RBD-V08H, the specific antibody S-D001, and the anti-β2-microglobulin antibody, were expressed and purified in-house.

### 5.2 Preparation of the CdS MetaSPR Chip

The MetaSPR biosensor was fabricated following the nanofabrication protocol established in our laboratory. A quartz substrate was patterned with a periodic tapered nanopillar array (top diameter: 100 nm, base diameter: 200 nm, height: 500 nm) using laser interference lithography to create a master mold. The mold was treated with a silane-based hydrophobic agent, after which a UV-curable adhesive was applied uniformly and covered with a PET film. The assembly was cured under UV light for 45 s at 105 mW cm^−2^ and then demolded, transferring the inverse nanocup array onto the PET substrate. Plasmonic metal layers (9 nm Ti, 15 nm Au, 25 nm CdS and 20 or 40 nm Au) were then deposited sequentially by electron-beam evaporation. The resulting sheet was cut into 13 × 8.5 cm sections and bonded to an open-bottom 96-well plate to form a 384-well biosensor plate.

### 5.3 Design and Simulation of the Nanocup Metasurface Chip

Numerical simulations were performed using the 3D finite-difference time-domain (FDTD) method with Lumerical FDTD software. Plane waves polarized along the x-axis were incident normal to the substrate (along-z). Perfectly matched layer (PML) boundaries were applied along z and x to suppress boundary reflections, with a periodic boundary along y. Reflection and transmission spectra of the two CdS chips were simulated in air and water to obtain the characteristic peak and valley wavelengths, their intensities and full widths at half maximum, from which the refractive-index sensitivity was evaluated.

### 5.4 Automated 384-throughput Imaging Platform and AI Image Analysis System

The optical responses of the CdS chips to varying sucrose concentrations were measured using the automated 384-throughput MetaSPR imaging platform to quantify sensitivity. Images were acquired at 100 Hz and processed by an encoder-decoder convolutional neural network (CNN) that performed well recognition, signal extraction and background correction.

For biological validation, the chip was washed and incubated with Protein A (ProA; 10 μg/mL in PBS, pH 7.4; 50 μL per well) at 4 °C for 14-16 h. The surface was blocked with 1% BSA at 37 °C for 30 min and washed with HEPES-ET buffer. IgG solutions of varying concentration (in HEPES-ET, pH 6.0) were injected into the ProA-modified wells and allowed to associate for 10 min, followed by dissociation for 15 min with 150 μL of HEPES-ET buffer.

### 5.5 Preparation of Protein A/G Biosensor Chips

The chips were cleaned and incubated with 50 μL of ProA or ProG (10 μg/mL in PBS, pH 7.4) per well at 4 °C for 16 h for passive adsorption. The surfaces were blocked with 1% BSA at 37 °C for 30 min to prevent non-specific binding. After removing the blocking solution, PBS containing 5% sucrose was added as a cryoprotectant to prevent protein degradation. Before use, the chips were stored under vacuum at 4 °C to limit metal oxidation.

### 5.6 Preparation of Tris-NTA Biosensor Chips

A metasurface chip (9 nm Ti, 15 nm Au, 25 nm CdS and 40 nm Au) was cleaned by alternating washes with anhydrous ethanol and deionized water, dried under N₂ and kept dry until use. EDC and NHS were dissolved in 50 mM MES buffer (pH 5.0) at 76.68 and 11.5 mg/mL, respectively, and combined at a 1:1 volume ratio. Thioglycolic acid (50 mM) was mixed with the EDC/NHS solution (1:1) and incubated at 37 °C with shaking at 300 rpm for 20 min. The photo-crosslinker was then added to the mixture and incubated at 37 °C in the dark for 6 h. Carboxymethyl chitosan (0.2 g/mL) was added and allowed to react for 10 min. The final volume ratio of EDC/NHS, thioglycolic acid, photo-crosslinker and carboxymethyl chitosan was 1:1:3:3. After vortexing, 5 μL of the mixture was spotted onto the chip surface and incubated at 4 °C for 16 h. The chip was washed, dried and blocked with 1% fish gelatin (100 μL per well) at 37 °C for 20 min, completing the first modification round.

For the second round, the chip was washed and shaken for 15 min. Tris-NTA (2.5 mM) and Na,Na-bis(carboxymethyl)-L-lysine (50 mM) were prepared in MES buffer (pH 5.0). Aliquots (3 μL) of each solution were spotted sequentially onto the chip and incubated at 4 °C for 14– 16 h. After additional blocking with 1% fish gelatin, the chip was washed and dried, yielding the Tris-NTA-functionalized MetaSPR chip.

### 5.7 Performance Validation of the Tris-NTA Biosensor

The optical responses of bare and Tris-NTA-modified chips to varying sucrose concentrations were measured. The chip was washed and activated with 50 mM NiCl₂ (50 μL per well) at 37 °C for 15 min, after which baselines were recorded. Three His-tagged proteins (RBD-V08H, FGFR1 and β2-microglobulin) were injected simultaneously, and association was recorded in real time for 10 min, followed by dissociation for 15 min (150 μL of buffer). The capture capacity of Tris-NTA was compared against conventional ANTA. Specificity was assessed by injecting S-D001 antibody at 20 μg/mL. Kinetics were measured by injecting S-D001 at varying concentrations over immobilized RBD-V08H (10 μg/mL).

Regeneration was performed with 150 μL of regeneration buffer (250 mM imidazole and 350 mM EDTA) under high-speed shaking for 5 min, repeated once (10 min in total). The wells were then washed twice with 150 μL of PBST (pH 7.4) for 1 min each. For subsequent cycles, activation with 50 mM NiCl_2_ (15 min) and ligand immobilization (5 μg/mL RBD-V08H, 20 min) were repeated.

### 5.8 Optimization of Tris-NTA Modification

To optimize the ligand dilution buffer, HEPES-ET and PBST at pH 6.0 and 7.4 were compared for their effects on RBD-V08H immobilization and subsequent S-D001 binding. For metal ion optimization, chips were activated with 5 mM CuCl_2_, 10 mM CoCl_2_, 10 mM ZnSO_4_, or 10 mM NiCl_2_ to evaluate their effects on ligand immobilization and analyte binding. Based on these results, NiCl_2_ and CuCl_2_ were selected for further concentration-gradient analysis. The PBS concentration of the NiCl_2_ buffer was then varied to maximize both ligand immobilization and analyte binding.

### 5.9 Sample Analysis

#### 5.9.1 Analysis of Purified Samples

Purified samples were analyzed using the Tris-NTA biosensor. All purified samples were diluted to 3.70 × 10^-7^ M in HEPES-ET buffer, and 50 μL per well was loaded onto the chip; antibody capture was monitored for 10 min. Dissociation was then initiated by adding 150 μL of HEPES-ET buffer to each well and recorded for 15 min.

#### 5.9.2 Detection of Samples from Different Sources

The antibody giving the highest Tris-NTA binding response in the preceding assays was used to detect antigens from different sources. Following the procedure described above, the purified antibody was first immobilized on a Tris-NTA biosensor. Human antigens were diluted to 3.78 × 10^-7^ M in HEPES-ET buffer and loaded at 50 μL per well. Binding was monitored for 10 min on the fully automated 384-throughput platform, and dissociation was recorded for 15 min with 150 μL of HEPES-ET buffer per well. Monkey antigens were measured identically, and all samples were cross-validated in parallel on a Biacore 8K.

Crude samples were analyzed on ProG sensors, first to establish how the dilution level of the primary-screen supernatants affects the measured affinity. Seven supernatants containing mouse IgG1 were selected, with IgG1 concentrations of 10, 20, 40, 50, 60, 80 and 100 μg/mL. These samples were diluted in PBST by different factors to reach the same final antibody concentration, and binding was measured for 30 min. In a second series, the same samples were diluted by identical factors and measured for 15 min.

#### 5.9.3 Assay of Crude Cell Supernatant Samples

Crude samples were analyzed using a Tris-NTA biosensor. Ninety-six crude cell supernatants were each diluted to 6.35 × 10^-8^ M, and 50 μL of each dilution was added per well in quadruplicate on a 384-well plate. Binding was recorded for 10 min on the fully automated 384-throughput MetaSPR platform. Dissociation was then followed for 15 min after adding 150 μL of HEPES-ET buffer to each well. The 15 samples giving the highest binding responses were selected for epitope binning.

#### 5.9.4 Epitope Identification

For epitope analysis on Tris-NTA sensors, the antigen was diluted to 10 μg/mL in HEPES-ET buffer (pH 7.4), and 50 μL was added to each detection well and incubated at 4 °C for 16 h. The 15 crude antibody samples were then diluted fourfold in HEPES-ET buffer (pH 7.4), and 50 μL of each was loaded into the detection wells of the MetaSPR sensor and monitored for 1 h.

In the tandem format on Tris-NTA sensors, the antigen was diluted to 10 μg/mL in HEPES-ET buffer (pH 7.4), and 50 μL was added to each detection well and incubated at 4 °C for 16 h. Antibodies from the different crude samples were diluted fourfold in HEPES-ET buffer (pH 7.4), and 50 μL was added to each well and monitored for 2 h until saturation was reached. Fourfold dilutions of the 15 crude samples were then added to the corresponding wells, and detection was continued for 1 h.

In the sandwich format on ProG sensors, antibodies from the different crude samples were diluted fourfold in HEPES-ET buffer (pH 7.4), and 50 μL of each dilution was added to separate wells on the MetaSPR sensor surface and monitored for 1.5 h. Each well was then blocked with 50 μL of blocking solution (fish gelatin and casein) at 37 °C for 20 min to suppress non-specific binding. The antigen was diluted to 10 μg/mL in HEPES-ET buffer (pH 7.4), 50 μL was added to each well, and the assay was continued until saturation was reached. Finally, 50 μL of each of the 15 crude samples, diluted fourfold, was added to the corresponding wells, and detection continued for 0.5 h. The entire process was monitored and recorded on the fully automated 384-throughput MetaSPR platform.

### 5.10 AI-Designed PD-L1 Binder Screening

Candidate PD-L1 binder sequences were collected from several generative models. Candidates were filtered on structure-based criteria: (i) contact with the PD-L1 core hotspots A56, A115 and A123; (ii) monomer pLDDT ≥ 80.0 and complex ipTM ≥ 0.75; (iii) instability index ≤ 40.0.

A composite score was then computed from interface contacts (40%), contact residues (30%), PRODIGY ΔG (20%) and hotspot hits (10%). The 48 top-ranked candidates were codon-optimized and transiently expressed in HEK293 cells. Crude culture supernatants were diluted and loaded directly onto the 384-throughput MetaSPR platform. Binding kinetics were measured at eight concentrations (48 binders × 8 concentrations = 384 channels) to derive the kinetic rate constants (k*_on_*, k*_off_*) and the equilibrium dissociation constants (K*_D_*). Top hits were cross-validated on a Biacore 8K.

### 5.11 Statistical Analysis

Data are presented as mean ± SD, n=3, unless otherwise stated. Statistical analyses were performed in GraphPad Prism 9.5.1. Group comparisons used two-tailed Student’s t-tests. Significance is denoted as *P < 0.05, **P < 0.01, ***P < 0.001 and ****P < 0.0001. Differences were considered significant at P < 0.05.

## Data Availability

All relevant data are available within the Article, Supplementary Information, Source Data file or available from the authors upon reasonable request.

## Code Availability

The custom analysis code used in this study is available from the corresponding authors upon reasonable request.

## Acknowledgements

This work was supported by grants from the National Science and Technology Major Project of China (2026ZD0128100, 2026ZD0128105), the Youth Program of the National Natural Science Foundation of China (32601989, 32471472), the Postdoctoral Fellowship Program of CPSF under Grant Number GZB20260937, the Shanghai Municipal Science and Technology Commission (25142201300).

## Author Contributions

**Liping Huang:** Visualization, Methodology, Data curation, Conceptualization. Methodology, Writing original draft, Resources, Funding acquisition. **Yuyan Wang**: Data curation, Visualization. Writing – original draft. **Yuxue Zhang:** Data curation. **Yannan Li:** Resources, Methodology. **Youqian Chen**: Visualization, Writing – review & editing, Resources, Funding acquisition. **Gang Logan Liu:** Writing – review & editing, Resources, Funding acquisition.

