## Supplementary Information for "Rapid Validation of AI-designed Drugs via 384-channel Parallel Kinetic Analysis in an Autonomous Laboratory"

*Corresponding author.

### Supplementary Information on the Results

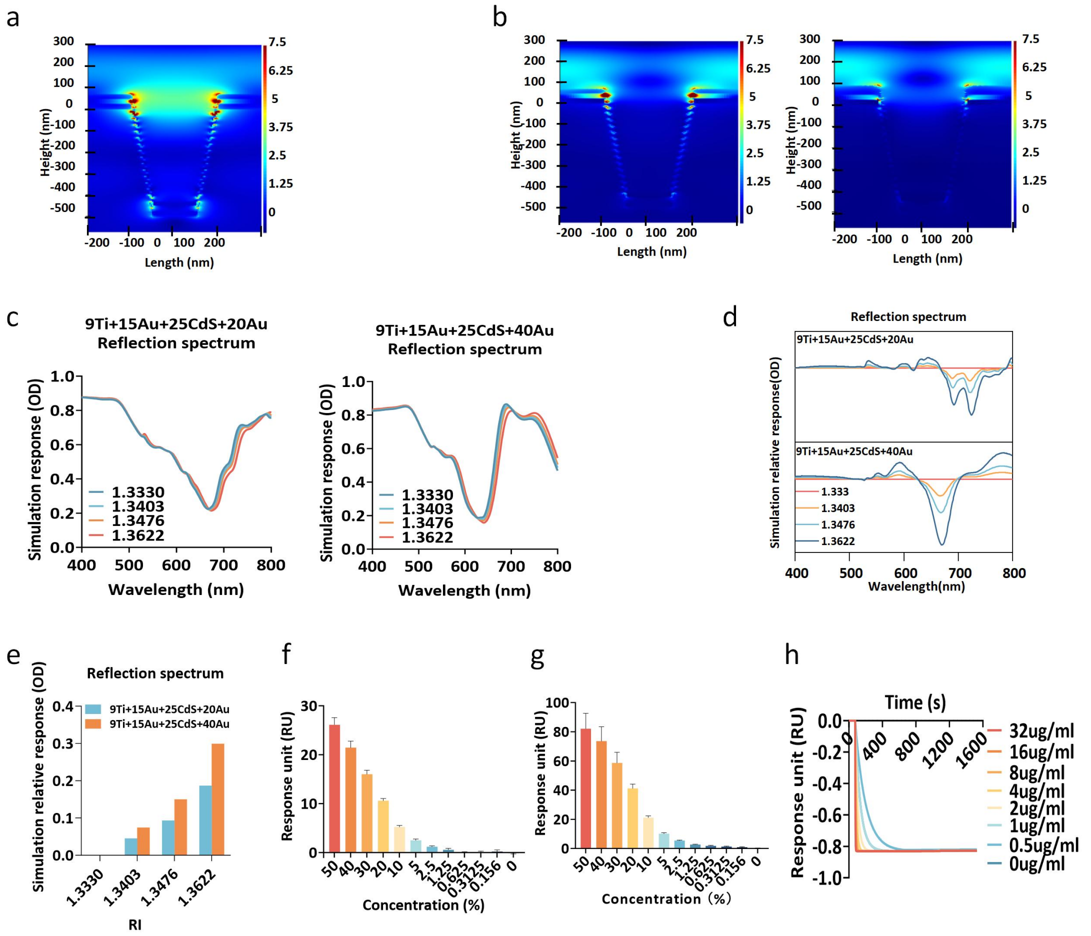


**Supplementary Fig. S1. Simulated transmission and reflection spectra.** (a) Finite-element simulation of the optimal electric field hotspots for the 9+15+25+20 gold chip at varied wavelengths. (b) Finite-element simulation of the electric-field distribution within the nanocup arrays of the 9+15+25+20 and 9+15+25+40 gold chips at the corresponding wavelengths. (c) Simulated reflectance of the gold chips of different thicknesses as a function of refractive index and wavelength. (d) Normalized reflectance spectra of chips with different thicknesses. (e) Comparison of differential reflectance spectra for SPR signal intensity (peak-to-valley) across chips of varying thicknesses. (f) Endpoint responses of the 20 nm Au chip to sucrose at different concentrations on the detection platform. (g) Endpoint responses of the 40 nm Au chip to sucrose at different concentrations on the detection platform. (h) ) Real-time kinetic sensorgram of Protein G-IgG binding measured on the 20 nm Au chip.

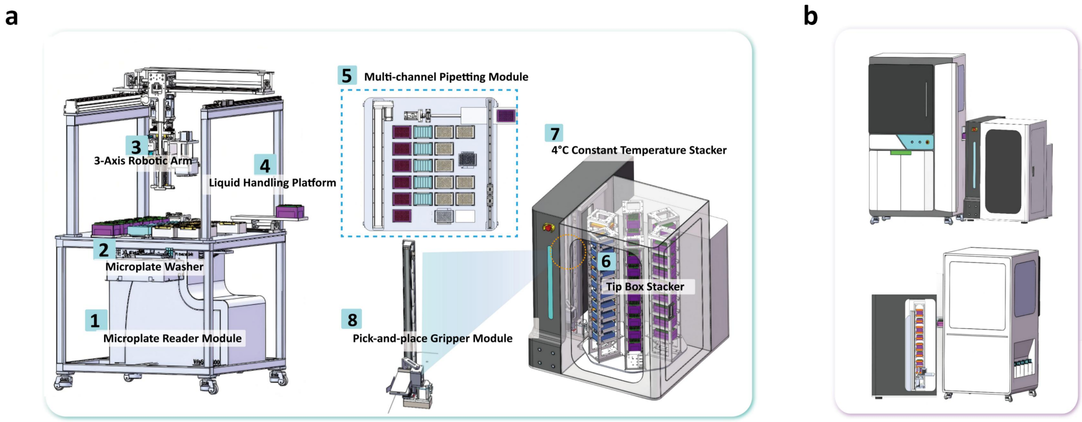


**Supplementary Fig. S2. Working principle and instrument architecture of the fully automated 384-channel high-throughput molecular interaction screening platform.** (a–b) Schematic diagram of the fully automated SPRM system hardware architecture: Cutaway view of the system, including front and rear perspectives. System components include a microplate reading module (1), a microplate washing module (2), a three-axis robotic arm (3), a liquid handling platform (4), a multichannel pipetting module (5), a tip stacker (6), a 4 °C temperature-controlled sample stacker module (7), and a sample handling gripper module (8).

**Supplementary Table S1. Spike-recovery assay**

| True concentration (μg/mL) | Slope | Back-calculated concentration (μg/mL) | Accuracy |
|---|---|---|---|
| 125.00μg/mL | 3.05 | 123.66μg/mL | 98.93% |
| 62.50μg/mL | 2.27 | 64.19μg/mL | 102.70% |
| 31.25μg/mL | 1.39 | 30.77μg/mL | 98.45% |
| 15.63μg/mL | 0.70 | 14.60μg/mL | 93.41% |
| 7.81μg/mL | 0.40 | 9.09μg/mL | 116.36% |
| 3.91μg/mL | 0.06 | 3.78μg/mL | 96.79% |
| 1.95μg/mL | -0.08 | 1.70μg/mL | 87.29% |
| 0.98μg/mL | -0.13 | 0.97μg/mL | 99.62% |
| 0.00μg/mL | -0.20 | — | |

**Optimization of assay conditions for Tris-NTA chip**

Several critical parameters govern the Tris-NTA modification process. The dilution buffer for the ligand (RBD-V08H) exerts a substantial impact on immobilization efficiency and subsequent molecular interaction assays. In this study, commonly used buffer systems including HEPES-ET (pH 6.0 and 7.4) and PBST (pH 6.0 and 7.4) were compared (Supplementary Fig. S3d). PBST at pH 7.4 gave the highest immobilization level of RBD-V08H and the strongest binding response to the S-D001 antibody among the four buffers tested, and was therefore used as the ligand dilution buffer in all subsequent experiments.

Tris-NTA is capable of forming stable six-membered ring complexes with metal ions such as $Ni^{2+}$ and $Cu^{2+}$ to establish a multi-site coordination binding system. The metal ion determines both the ligand immobilization efficiency and the performance of the subsequent interaction measurement. We systematically evaluated the activation performance of four metal ions: $Ni^{2+}$, $Co^{2+}$, $Zn^{2+}$ and $Cu^{2+}$ (Supplementary Fig. S3e). $Ni^{2+}$-activated chips achieved the highest immobilization capacity of RBD-V08H, whereas $Cu^{2+}$ activation produced stronger binding responses between the ligand and analyte. The two stages therefore favour different metal ions. Further concentration-gradient assays revealed that $Cu^{2+}$-mediated binding responses were substantially greater than those of $Ni^{2+}$ at elevated analyte concentrations, suggesting that $Cu^{2+}$ is more suitable for interaction assays with high-concentration analytes (Supplementary Fig. S3f).

To further optimize metal ion activation conditions, we assessed the effects of dilution buffers containing varying PBS concentrations on $Ni^{2+}$ activation and downstream detection (Supplementary Fig. S3g,h). $Ni^{2+}$ diluted in high-concentration PBS significantly promoted ligand immobilization and ligand–analyte binding. Phosphate in PBS may complex with the metal ions and increase their coordination to Tris-NTA, which would account for the higher responses.

Finally, the regeneration protocol described in Methods was applied to validate the regeneration performance of the Tris-NTA chip. After six regeneration cycles, the binding responses toward RBD-V08H and kinetic sensorgrams remained highly consistent without obvious signal decay. The interface therefore tolerated at least six regeneration cycles, which is the prerequisite for repeated kinetic measurements on a single chip.

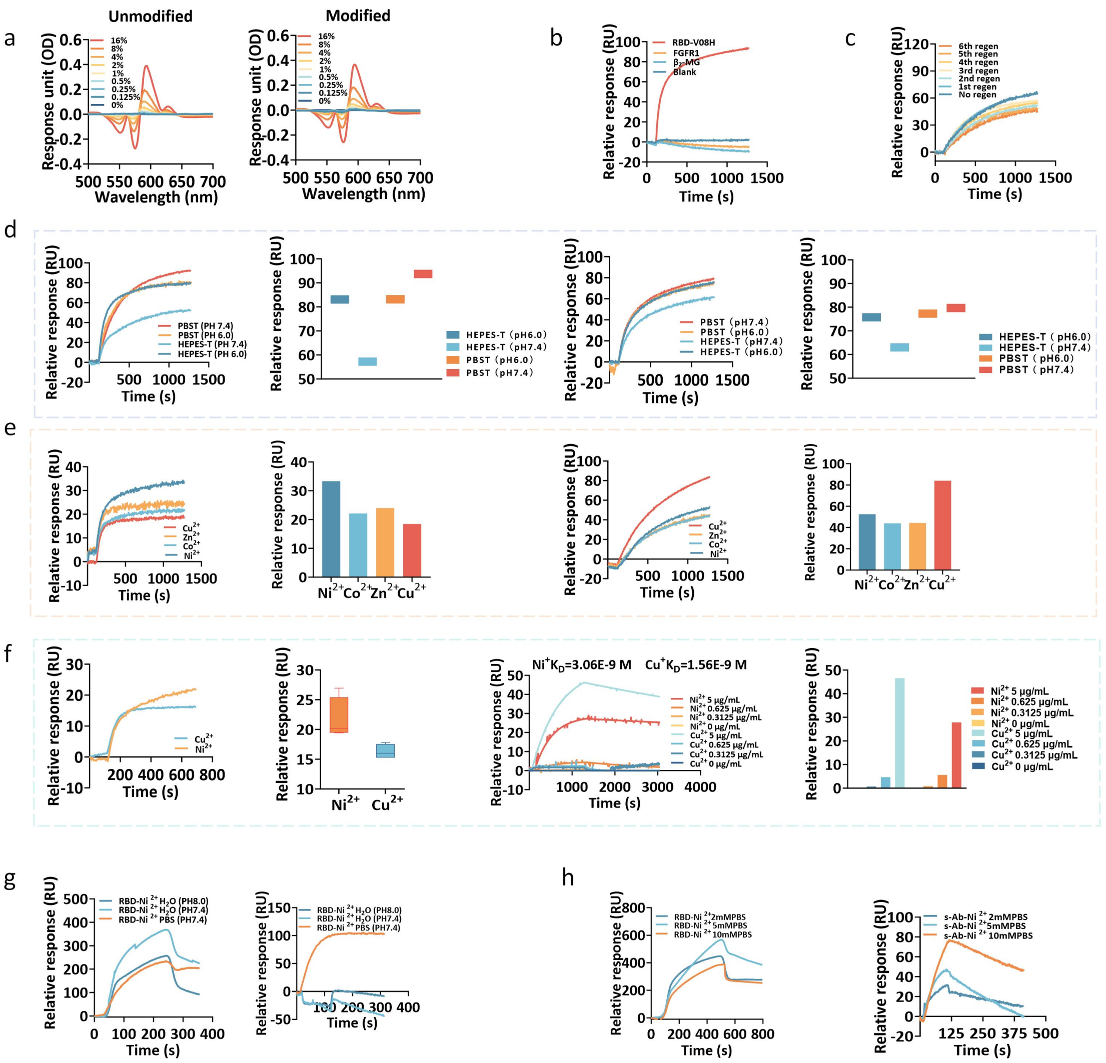


**Supplementary Fig. S3. Optimization of assay conditions for Tris-NTA chip.** (a) Reflectance spectra of the unmodified and modified biosensor surfaces. (b) Specificity test of the Tris-NTA biosensor. (c) Real-time sensorgrams of Tris-NTA chip during regeneration cycles. (d) Real-time sensorgrams and endpoint comparison for ligand (RBD-V08H) immobilization in different dilution buffers, and for the subsequent binding of the analyte (S-D001 antibody). (e) Real-time sensorgrams and endpoint comparison for ligand (RBD-V08H) immobilization after activation by different metal ions, and for the subsequent binding of the analyte (S-D001 antibody). (f) Real-time sensorgrams and endpoint comparison for ligand (RBD-V08H) activated by $Cu^{2+}$ versus $Ni^{2+}$, and for the binding of the analyte (S-D001 antibody) at different concentrations. (g) Ligand and analyte binding with $Ni^{2+}$ diluted in various buffers. (h) Ligand and analyte binding after $Ni^{2+}$ activation using PBS of different concentrations.

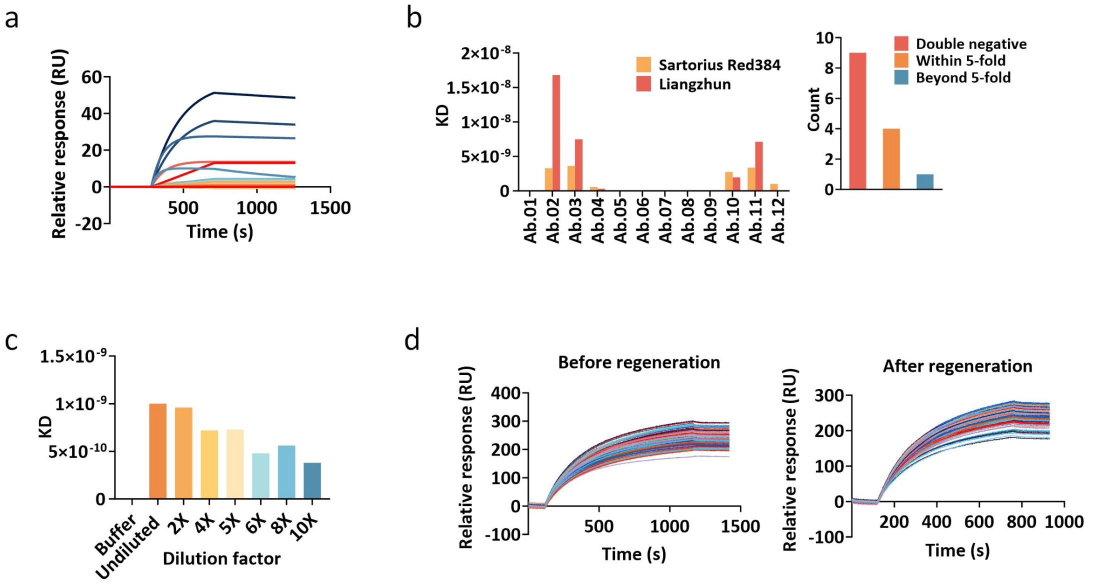


**Supplementary Fig. S4. Detection of samples from different sources.** (a) Real-time sensorgrams for affinity measurement of purified antibody. (b) Comparison of $K_D$ values obtained by MetaSPR and Sartorius Red384, and the count of $K_D$ pairs within a 5-fold difference. (c) KD values determined for mouse IgG1 samples at different dilution factors in PBST. (d) Real-time sensorgrams for consistency validation of the Protein G chip before and after regeneration.

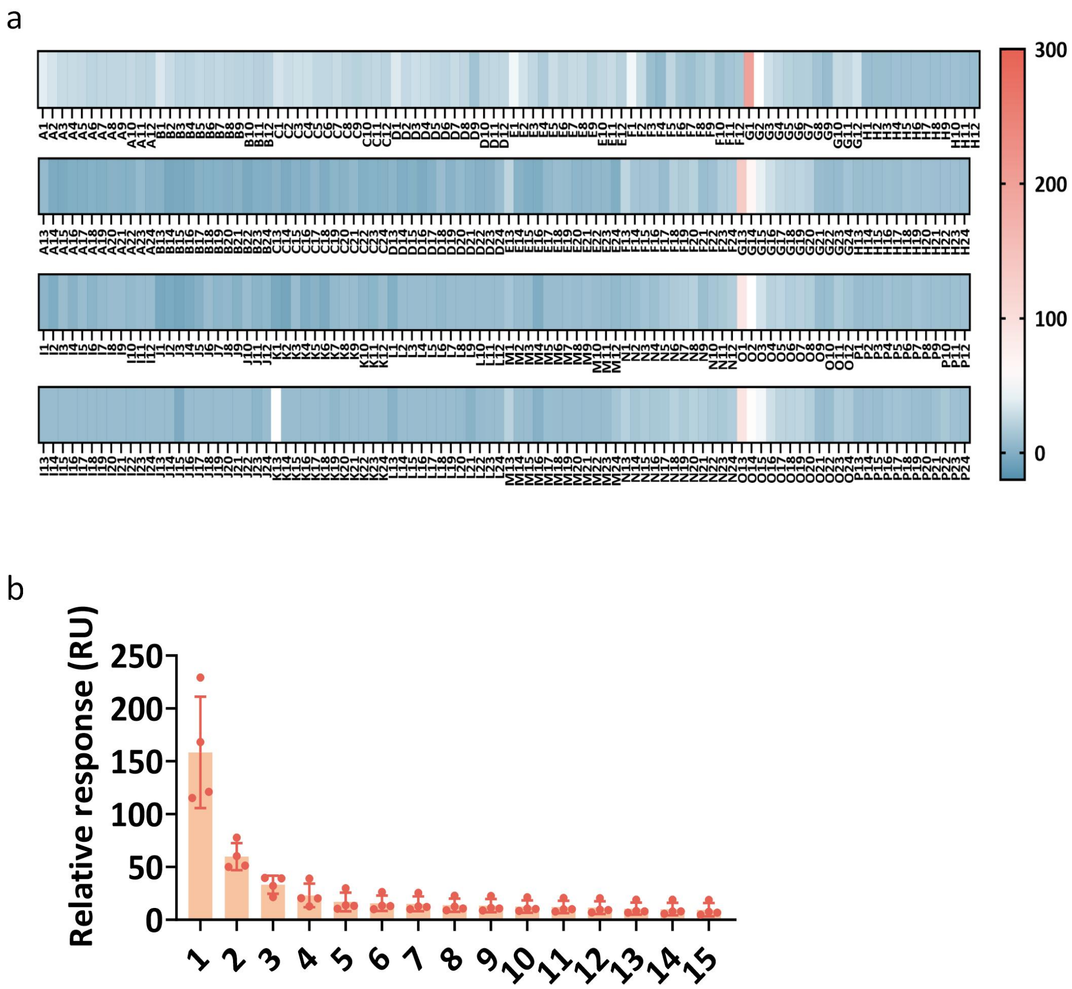


**Supplementary Fig. S5. Consistency of 384 channels.** (a) Heatmap showing endpoint responses of real-time kinetic signals for cell supernatant samples, synchronously collected from 384 sensing channels in one assay. (b) Endpoint responses of the 15 highest-ranked candidates across repeated screens.

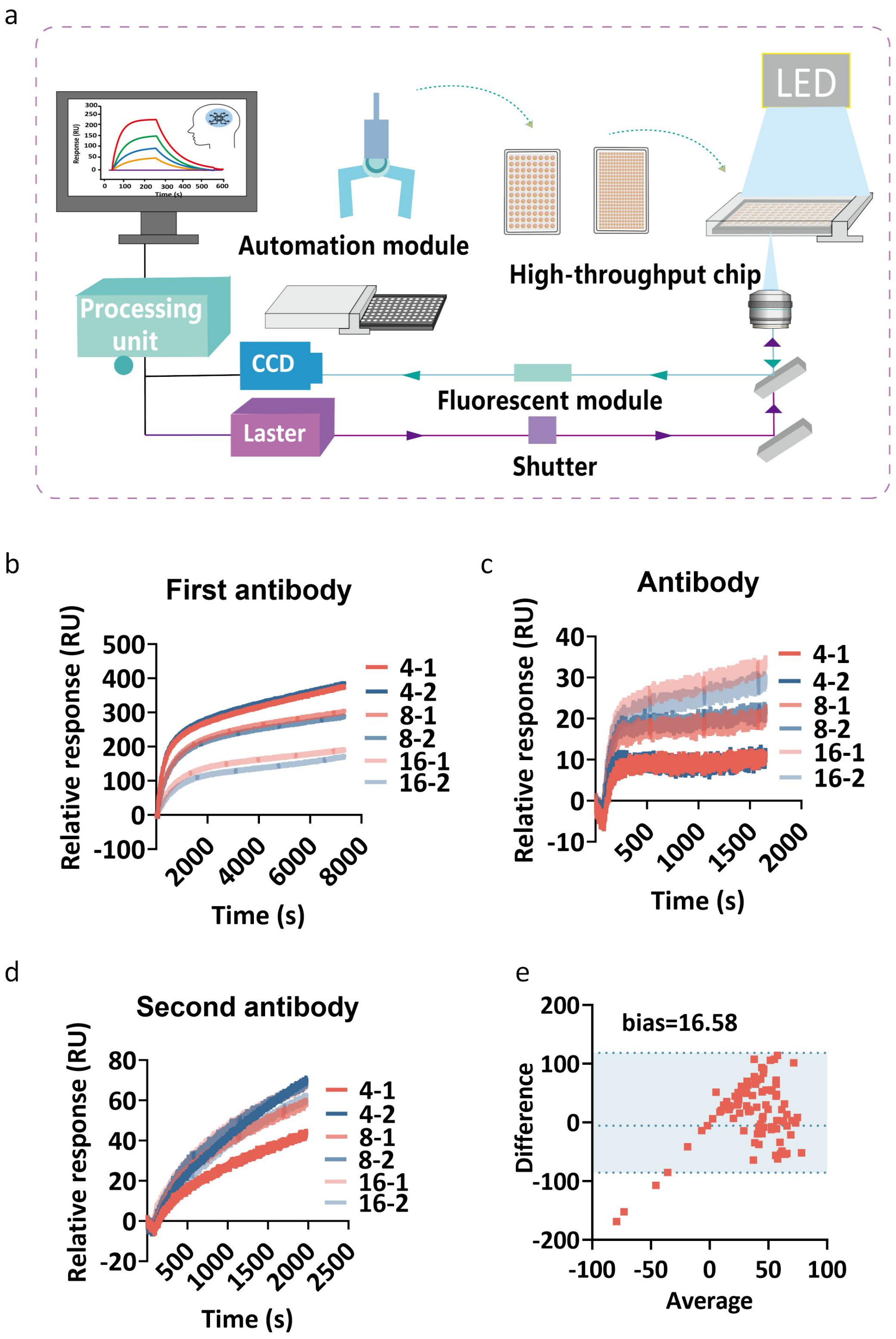


**Supplementary Fig. S6. Optimization of epitope pairing conditions.** (a) Schematic illustration of the hardware detection workflow. (b-d) Epitope binding tested in the sandwich format at four-, eight- and 16-fold sample dilution; the fourfold dilution was optimal. (e) Bland–Altman plot assessing agreement between the tandem and sandwich formats.

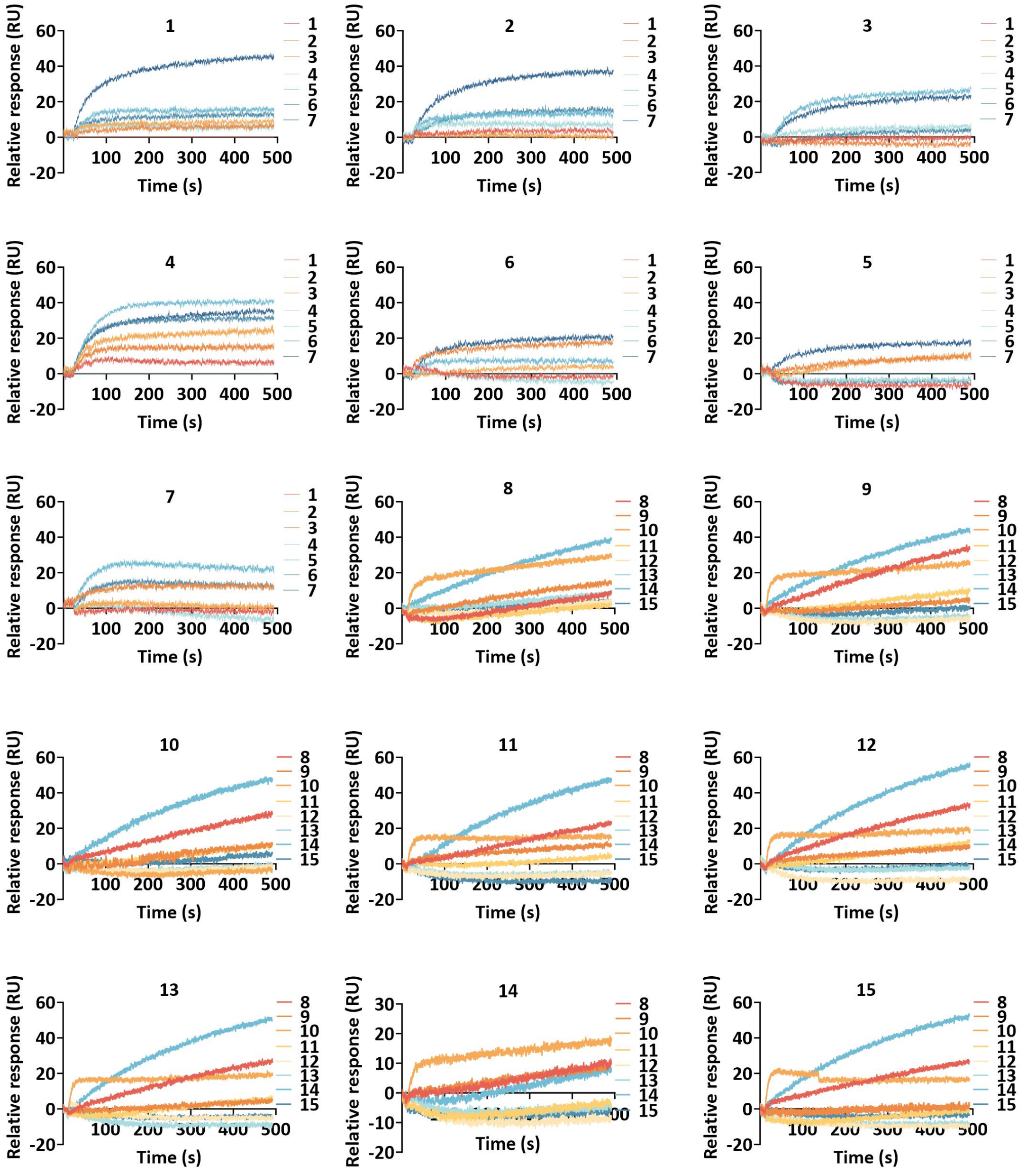


**Supplementary Fig. S7. Real-time sensorgrams of secondary antibody binding in the tandem assay.** Samples 1–7 are grouped for epitope pairing, and samples 8–15 are grouped for epitope pairing.

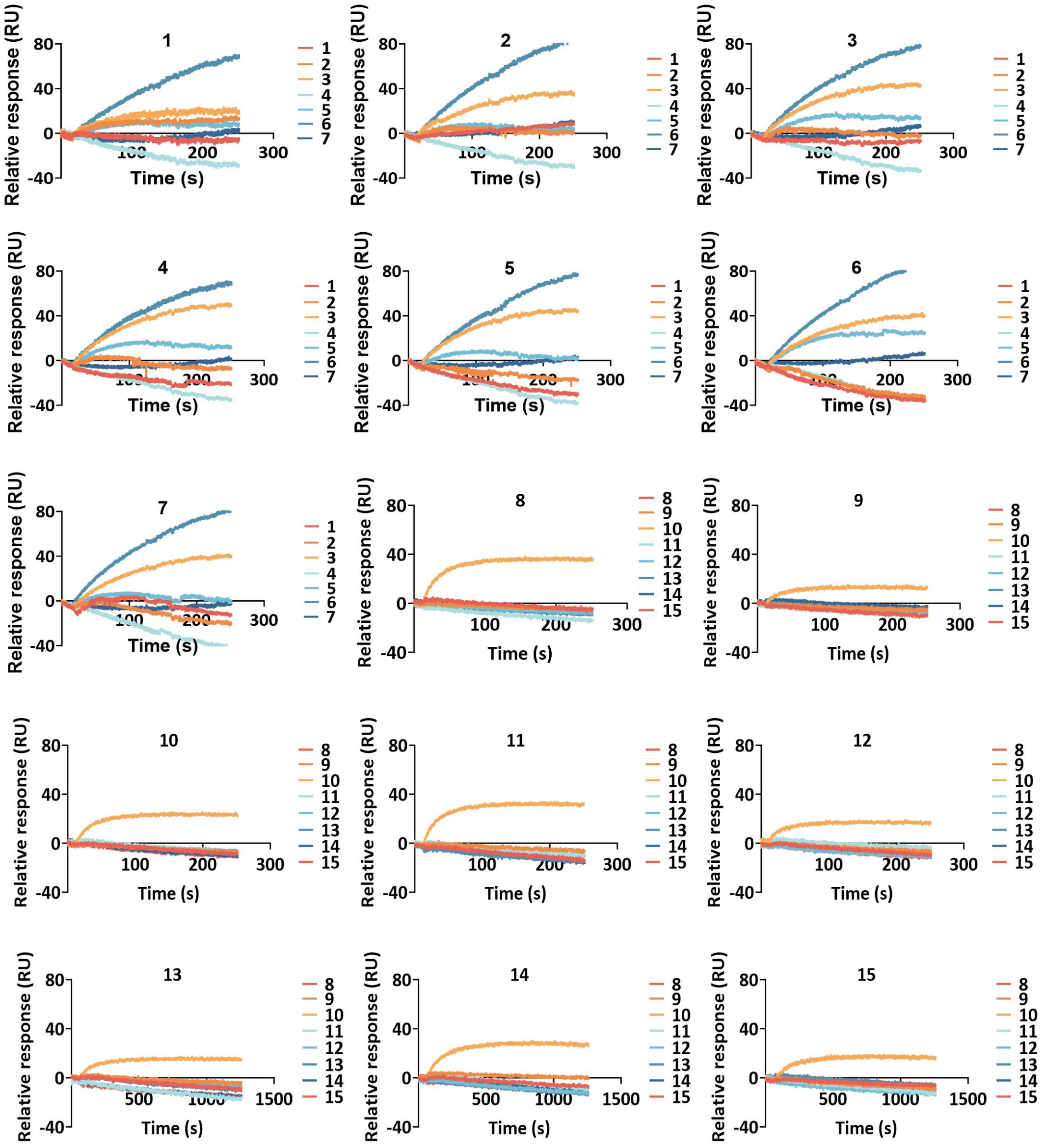

**Supplementary Fig. S8. Real-time sensorgrams of secondary antibody binding in the sandwich assay.**

a

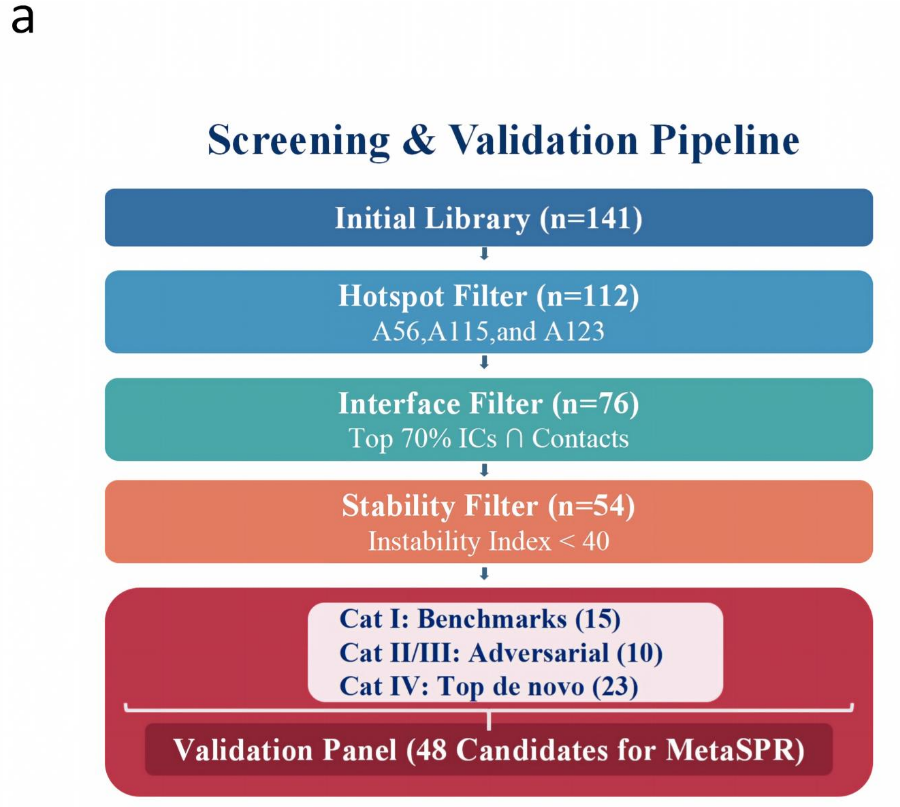


**Supplementary Fig. S9.** Screening strategy and composition of the 48-candidate test panel. The three-tier convergence workflow applied to the 141 de novo designed sequences is shown: binding-hotspot constraint, interfacial atomic-contact evaluation and sequence-expression stability. The composition of the final 48-candidate panel, which comprises four functional subgroups (Cat I–IV), is also shown.

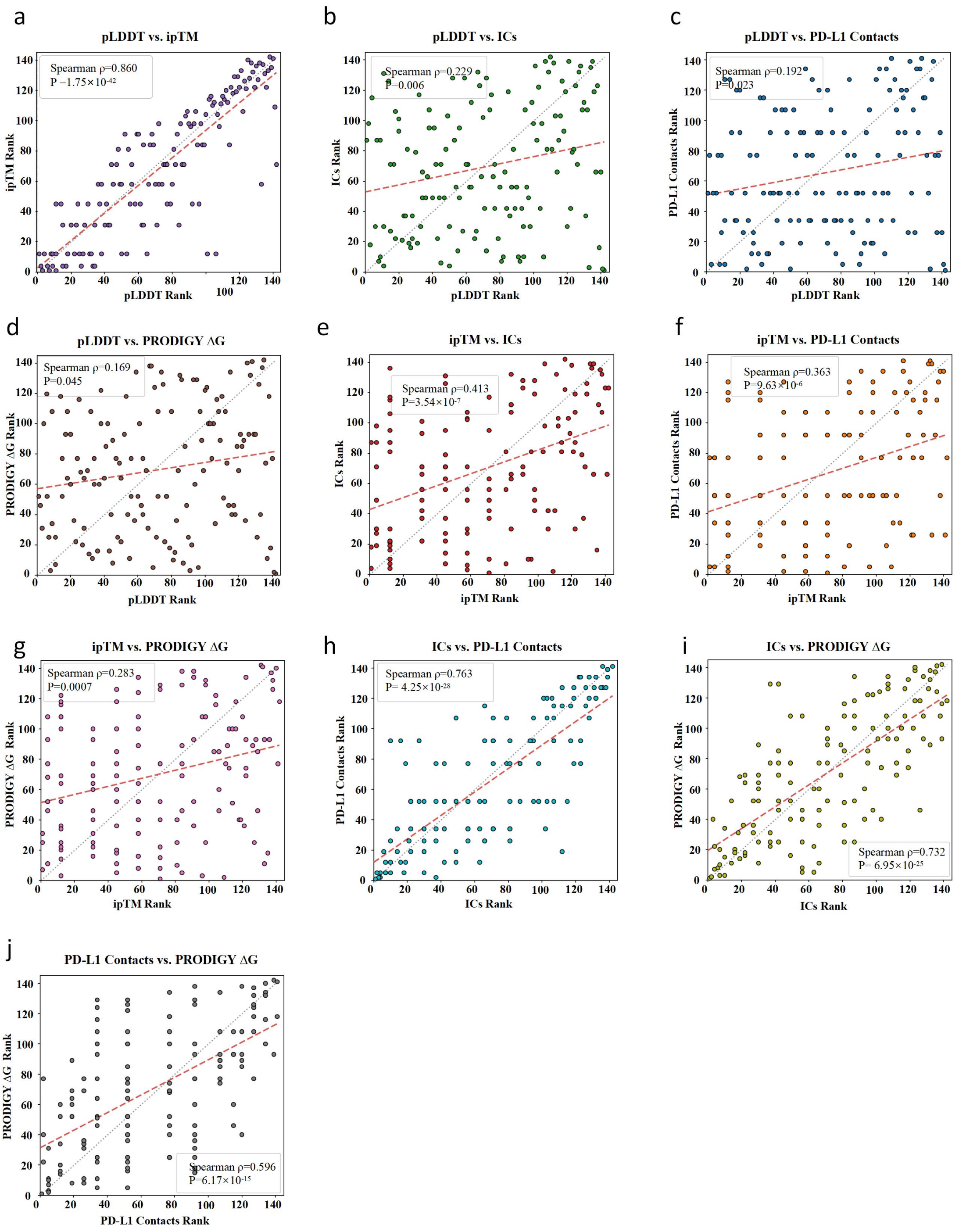


**Supplementary Fig. S10. Overview of dispersion and correlation across all ten core metrics.** AI-derived metrics and physical contact metrics exhibit evident cross-domain decoupling. n = 141; both axes show rank values (1 = best; range 0–141). The grey dashed line denotes y = x; the red dashed line denotes the linear regression fit. Strong intra-domain correlation: (a) pLDDT versus ipTM: ρ = 0.860, p = $1.75\times10^{-42}$, slope = 0.91. Points cluster tightly around the diagonal, indicating high concordance between the monomer and complex confidence scores of AlphaFold2 (AF2). In contrast, the cross-domain pairs (b–g) were widely dispersed (slopes < 0.4), indicating that high-confidence structures do not necessarily form large contact interfaces or favourable binding energies. (b) pLDDT versus ICs: $\rho = 0.229$, $p = 0.006$, slope = 0.23. (c) pLDDT versus PD-L1 Contacts: $\rho = 0.192$, $p = 0.023$, slope = 0.21. (d) pLDDT versus PRODIGY ΔG: $\rho = 0.169$, $p = 0.045$, slope = 0.17. (e) ipTM versus ICs: $\rho = 0.413$, $p = 3.54\times10^{-7}$, slope = 0.39. (f) ipTM versus PD-L1 Contacts: $\rho = 0.363$, $p = 9.63\times10^{-6}$, slope = 0.36. (g) ipTM versus PRODIGY ΔG: $\rho = 0.283$, $p = 0.0007$, slope = 0.28. High concordance was also observed among the physical interface metrics: (h) ICs versus PD-L1 Contacts: $\rho = 0.763$, $p = 4.25\times10^{-28}$, slope = 0.77. (i) ICs versus PRODIGY ΔG: $\rho = 0.732$, $p = 6.95\times10^{-25}$, slope = 0.72. (j) PD-L1 Contacts versus PRODIGY ΔG: $\rho = 0.596$, $p = 6.17\times10^{-15}$, slope = 0.58.

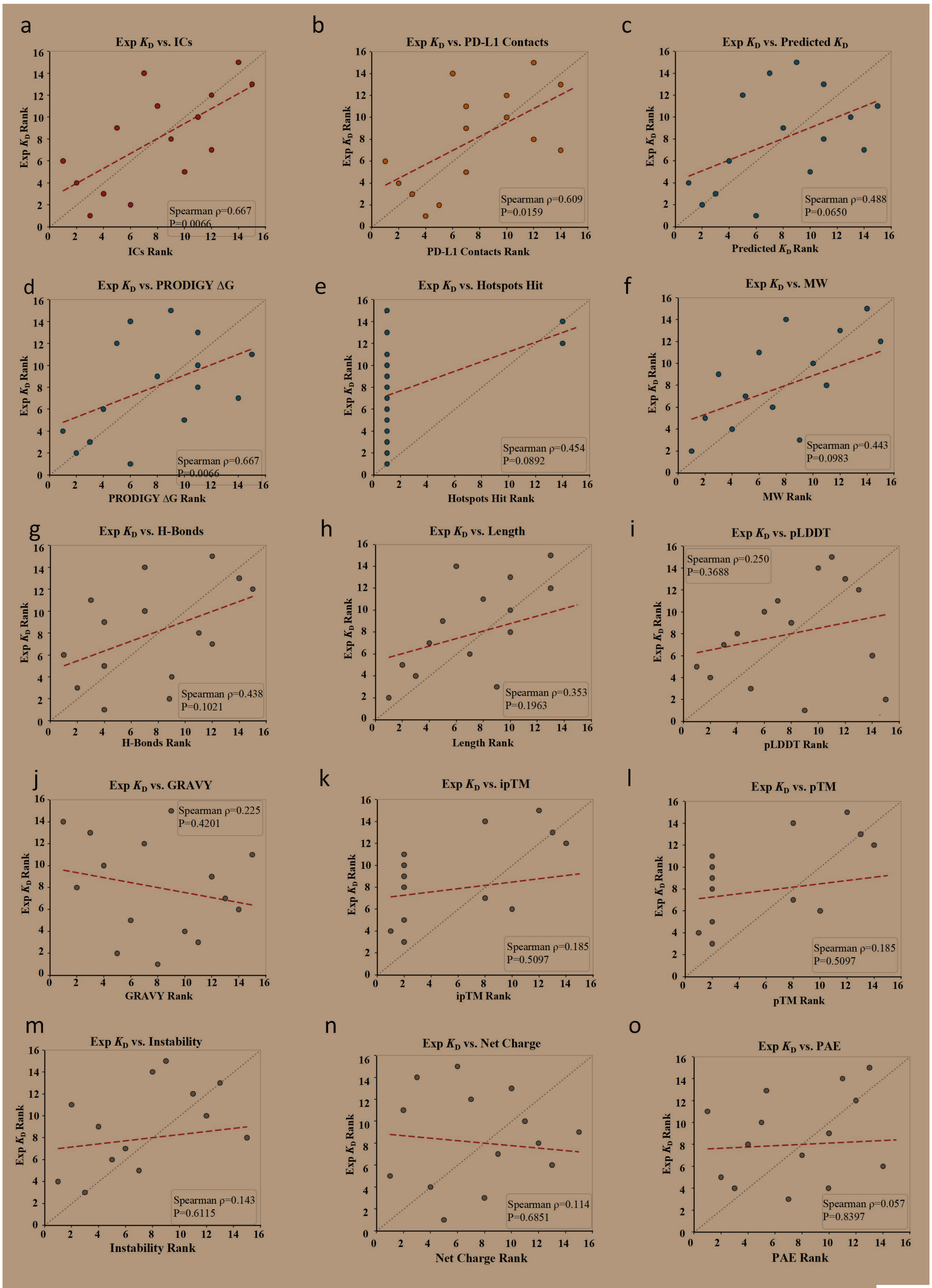


**Supplementary Fig. S11. Independent performance of individual metrics** for experimental binding affinity prediction, validated using 15 literature benchmark entries with reported $K_D$ values ranging from 0.14 nM to 2 µM. n = 15 literature benchmarks. The y-axis shows the rank of the experimental $K_D$ (1 = best); the x-axis shows the rank of the corresponding computational metric (1 = best). Physical and binding-energy metrics (strong predictive power): (a) Experimental $K_D$ (Exp $K_D$)

versus interface contacts (ICs): ρ = 0.667, p = 0.0066, slope = 0.68, the highest correlation among the metrics tested. (b) Exp $K_D$ vs. PD-L1 Contact Residues: ρ = 0.609, p = 0.0159, slope = 0.64; greater residue contacts correspond to stronger binding affinity. Metrics with moderate correlative trends: (c) Exp $K_D$ vs. PRODIGY Predicted KD: ρ = 0.488, p = 0.0650, slope = 0.49. (d) Exp $K_D$ vs. PRODIGY ΔG: ρ = 0.468, p = 0.0784, slope = 0.48. (e) Exp $K_D$ vs. Core Hotspots Hit: ρ = 0.454, p = 0.0892, slope = 0.44. (f) Exp $K_D$ vs. molecular weight (MW): ρ = 0.443, p = 0.0983, slope = 0.44. (g) Exp $K_D$ vs. number of interfacial hydrogen bonds: ρ = 0.438, p = 0.1021, slope = 0.45. Metrics with no significant correlation (AI confidence scores and physicochemical descriptors): (h) Exp $K_D$ vs. sequence length: ρ = 0.353, p = 0.1963. (i) Exp $K_D$ vs. pLDDT: ρ = 0.250, p = 0.3688. (j) Exp $K_D$ vs. GRAVY (hydropathy): ρ = −0.225, p = 0.4201. (k) Exp $K_D$ vs. ipTM: ρ = 0.185, p = 0.5097. (l) Exp $K_D$ vs. pTM: ρ = 0.185, p = 0.5097. (m) Exp $K_D$ vs. instability index: ρ = 0.143, p = 0.6115. (n) Exp $K_D$ vs. net charge: ρ = −0.114, p = 0.6851. (o) Exp $K_D$ vs. mean PAE: ρ = 0.057, p = 0.8397.

**Supplementary Table S2. Amino-acid sequences of the 48 candidate PD-L1 binders.**

| No. | Sequence |
|---|---|
| 1 | SAEEKILANLEAMKAKALAAKTEEEKLFYAKALLAVAISYAIRGDYELARRAAEL AVEVIKSLSKEEQKKVMDFLINIIKNITDPEDREKAIELAIAIAERLDEEVREEALKK IEELKKE |
| 2 | EEDCKVHCVKEWMAGKACAERQKSYTIGRAHCSGQKFDVWKCLDHCAAP |
| 3 | SKAEAAANRMKRFLDGLKISIPELRDLIEKYGEKIVEAIKAGDKEKALKYAEELAK KIKEVLTDDPVFAENLAKFVIVYVESLLEEL |
| 4 | LKEEALELADEVIKLAEELGWKDHVKAVEALKEAVEKSTDERFLASAKAFLEVL KEVLLEEKKA |
| 5 | MYEVVIEGEKSVAEFIKLIAEQLGAEAEVEGDKAVIRTERREDAERLAEAAKRFG AEAEVRE |
| 6 | AAAESAALAAALAALRKTEVGPFARQYIIYRLLAQVAEAKGDEETAKRAKAKAE ELAKAYPEAAAAADAALALAQAALAA |
| 7 | AQTILLRLEGMDSTSSASSIERAIAKVPGVQQCIVFFEMNLAIVTYHGETTPQILTD AVERAGYHARVS |
| 8 | NLLRSYENSFKIQLILAKLELAHAPSQPLSQRNEELKRVEQRQDRLFDLLDQMDVE VNNSIGDASERATYKAKLREWKKTIQSDIKRPLQSLVDSGD |
| 9 | SESEEWIKEITDAANQAVALAKAGKTAEAYALLDATAAKAAAKSPSRGALAKGV ADAMRVYIDKYAAKKA |
| 10 | NLLTSYEGSFKIQLILAKLELAKAPSQPLSQRNEELKRVEQRQDRLFDLLDQMDVE VNNSIGDASERATYKAKLREWKKTIQSDIKRPLQSLVDSGD |
| 11 | SNEEKAEDLRFMARAVLGVGATASAAETAAFLNDVAAAAARLDPEAGAAVAAI GAAAAASGDAKAAAKAAAAVLEAAAEEFL |
| 12 | SSIESMKWSMIRQQILCQLETGIDQQKANDVIEGNIDVEDKKVQLYCECILKQFHIL DKNNVFKPQGIKAVMELLIDENSVKQLVSDCSTISEENPHLKASKLMQCISKYKT WKSFDFL |
| 13 | NLLESYEWSFKVQLILAKLELAKAPSQPLSQRNEELKRVEQRQDRLFDLLDQMDV EVNNSIGDASERATYKAKLREWKKTIQSDIKRPLQSLVDSGD |
| 14 | VEEIMEDPEKTYEKLWKELKEKLKGKSPHEIRTAFHEKFWEWIDEVIAKKLPNEAI IKLLWVVRKFDDEMEKLEEEEKA |
| 15 | SSIESMKWSMIVQQILCQLETGIDQQKANDVIEGNIDVEDKKVQLYCECILKQFHI LDKNNVFKPQGIKAVMELLIDENSVKQLVSDCSTISEENPHLKASKLMQCISKYKT |

| | |
|---|---|
| | WKSFDFL |
| 16 | SLEEAADEAAVAVVLAMQTVSKAGAEEAAEIRKAADDAADAIRKNLDADFAAE LAAELRKAADEAVALAKAGRVEEARRHFAEVATEVIKRAREYAEKA |
| 17 | LEEERREFEEVLERLLEVAEEDPEVRERAEEMVRRAREVAEREGVAAGAAEAAV GALRIASEASEEVRKRYLELASELQAIGQRVVRA |
| 18 | SSPEEVIKRLLEEAVKRAAEAAAHIARTDPEAAAALRAAAEKAKADPKHAGDLA ALARELAERFAADPEARRLLRGVASIADMITLILEDK |
| 19 | EEEEKKEKANLLRVEAAQKLSLVAEALPEVREKALEAIRKIFDKETIEEGVELAREI AERVAAEGTEEQARLAREAVEKAEEALKLEK |
| 20 | REEKIEKVREAAEEAKKAGAIEDYEIEGDEVKFYRKDWKPESLLIVMKAAQVAAL EGFKPTIVKS |
| 21 | AQTILLRLEGMDCTSCASSIERAIAKVPGVQQCIVFFEMNLAIVTYHGETTPQILTD AVERAGYHARVS |
| 22 | SMERIISDIRFIAGYLYAVYSHEARHNPENEVAKEIADFMYAVKTNPLEKWPEFAK KLSEESKKIIYDVFKSIGEQHNNSEMIEIVEKIL |
| 23 | EEEEEEKKKEEKEKEKKKLLKEAAKELEDIYKNVDEFSIDFMSAVLEQIGERLAK YDEEIAKFMKEAGERLKTATKEEAKEIIKEAIKKLKEAAK |
| 24 | KREEALRKAAEAAVKAMEMAKKASDEAKVEIMEHFSRAMRAIKEGDFEEAIRHL RKAAEVAEEKVGKEVGEFIREAAREVERIA |
| 25 | SDELRVEAVQVAQEASVAIARAERAVAEGDAEEIRRAAEELERAARRAEEQAERL EEAGDKGMAEFLRRVVERAREAAARARRAAEELE |
| 26 | DEFLERVRELAERLREAAEGTDEEARAVTEEAVRELEEIAEELLKDPKTRVAGILA RAAARQTRALLARSDDARAVARQAAANAELVAMRLEDI |
| 27 | MSKRDELLTKALEYLQKATKLVYEHDPETAREIAHHFSKAVADPEKAEEELEKAL ELAEKAAAEMERGEELRELMEKAKEYVLEYVKLKK |
| 28 | MEEVKEFLAKNSPYSKNFLDDFIEKHREGKVTEEDWEKLKDYFINAAKLVASEDE TFIKHWEKDVEQMMEWMKTATPSDFEFVVRFYLSVLQKMLEMK |
| 29 | SVVDRVREALGHTFRTRFGGREIVVFLGTEDDSLWIMVVYPDGRLKVFRFRDLAG LVESGEFDRLLELLRSRDLEAFEAFLAS |
| 30 | SEEKKKLALEAAAKLSQAMKVLAEKRDDVKAAAEEAVKLMKEAAKLAAKADE ELKKKVEEALKEVEKNRDDAEKMMEIVAKAIVDVMKYISKLD |
| 31 | SPAERIRELAREARRAVEAGDGERAARLAQEMFRLALEDGEVRPLALKELLEVM RLAAERDPETARLIAEAAAEVDPRHAHFLTSYVDQLVERVAA |
| 32 | SAAAAARARTAAANAAALAKKMLEAGVSPENARALVEAAAARLTPEAGAALRA EGEALVAEYAARREAEAA |

| | |
|---|---|
| 33 | AKREEIQKIIEELLKEAKEFLKEAKGHPEEPLIKEAIKLLEEAKKLLEAGREKEAIV KAIVAADKLIAALKKYDPSKKERIEKIKEKRKRLKELFE |
| 34 | SIEEMVEEMEKLLEEAAKIVPEIKELLEKAKEIAKKAIEIAKKKGNGEAVAKIFRD MAEMTLRAAKFDKKGAENAALLMIKRAEDIAE |
| 35 | LDEFVRFARSAVEFIALMIKEGVPESILDTTIEIFKERISELIREYGAEAGLSAEEREA FAKEFAAALDGMKEDPEAYRERALALVEEFARKAEEAL |
| 36 | SKEEKIKEIEKKIEEATKLMVEGIREGDKEKVEKGKELMVKAAKEVGDEATAEFIE SFAEKAIRVAEEDKEKAVKMVKEGGKLVKVRLKDRIEE |
| 37 | GEVYERIVKVTYTFTPDGKMIVTFEVIDENGNVIPDEPVPPVIEEDMENVDPNDSIV LNFLKFRSIHEYEYYTGEKVLSYEYTIEKSYDPEKGEIITFTFKLTVER |
| 38 | DLEELAEEVAEKLAKIGDEELAKEFGEELKKALEIVGDEEIKEQFLKAVDLLENGT DEEALLTLMEAASQAVLKNPAAAPYAKKMVTLFRQMIKLKR |
| 39 | SKKERAVKQLKEKKEKFKEIADELLAEAKADPSKAAEAAKTIDELIKMVKLMAK DKAEETGAEEELNKIAEEFEKYAKEVRAEVEAAGAAAAA |
| 40 | SESEEWIKEITDAANQAVALAKAGKTAEAYALLDATAAKAAAKSPSRGALAKGV ADRMRVYIDKYAAKKA |
| 41 | DKESAAKKAIQLLEIARERVVADPAAAKAFKEAAKLVEVMLKDAGADEEKAEK VEEKAEKAADAYAAGDKETADKYYEEAVELVKEAA |
| 42 | DEMDKLKVEAMKYGSKAIKIASEKDEELAERAAEALSLAMTDPEKGAKELRKIA EELEEKGLPEAAEYLRKAADYFERYAKLLKE |
| 43 | EARELVKEAVEKFEKAAEEALKDFEKAEKLAKEALEKLYEAAKLDPSLRPVYQE AALHLVKALAHDDDPEKASEELKKAAEVAKKELEK |
| 44 | AEEKAREELEKARKLAEEVKKKHPALGEALLSMLDLAEELFERDREAARRLLRA LAAFAEAAAKALAEEDEEAAKLVKELAEKVAEAARLLE |
| 45 | MSQVEKEAEALVEAGRYVAERAAAEVGDRERARELVEGFADFLALLVKENPKR YKGAVEAAEAAAEVLAAADAEFGEEARKLMFEKIEKGKAEAEAK |
| 46 | EKLKEKMKKFVEDIAYLVGQAVYAETGDKEKAKEEAEKIKKEFEKALEDIEKAL KFLEELKEELREKAKEFGRAGSEIFKLAALKIEDFLEEYRK |
| 47 | DARVQVVVELLEKAAEEVARLDPSVAEKVKKALTEAAKEAEKMSAKEAAEYAA KVAEELLKELADKLSPEAKEVLQGAVTFLTKLAEIA |
| 48 | GAAEVEKYRKKMIELFEKFKDYFLSITEEGSELHKKFLKEYKKILKLLEEGKYKEA MEKIYVLVFDAMDKKSEHREELFMMLAEINWYYFQMMKAE |
